# Dynamic hotspots in the Uba7 UFD direct UbcH8 recognition

**DOI:** 10.1101/2025.06.14.659664

**Authors:** Çağdaş Dağ, Mahil Lambert, Alp E. Kazar, Kerem Kahraman, Oktay Göcenler, Woonghee Lee, Cansu D. Tozkoparan Ceylan, Frank Löhr, Jin Gon Shim, Arthur L. Haas, Volker Dötsch, Joshua J. Ziarek, Emine Sonay Elgin

**Affiliations:** Nanofabrication and Nanocharacterization Center for Scientific and Technological Advanced Research (n^2^STAR), Koç University, İstanbul, Turkiye; Koç University Isbank Center for Infectious Diseases (KUISCID), Koç University, Istanbul, Turkiye; Department of Chemistry, College of Sciences, Muğla Sıtkı Koçman University, Muğla, 48000, Turkiye; Centre for Biomolecular Magnetic Resonance, Institute for Biophysical Chemistry, Goethe-University of Frankfurt/Main, Germany; Department of Chemistry, University of Colorado Denver, Denver, CO 80204, USA; Department of Pharmacology, Feinberg School of Medicine, Northwestern University, 320 East Superior Avenue, Chicago, IL, 460611, USA; Department of Biochemistry and Molecular Biology, LSUHSC-School of Medicine, 1901 Perdido Street, New Orleans, LA, 70112, USA

## Abstract

The ISGylation pathway, a key post-translational modification, plays a pivotal role in the innate immune response by covalently attaching ISG15 to target proteins. This cascade involves a series of enzymatic steps, including activation by the E1 enzyme Uba7, conjugation by the E2 enzyme UbcH8, and ligation by an E3 ligase. Central to this process is the ubiquitin-fold domain (UFD) of Uba7, which facilitates the transfer of ISG15 to UbcH8. However, the structural and mechanistic details of this interaction remain poorly understood. In this work, we present the solution NMR structure and functional analysis of the ubiquitin-fold domain of human Uba7, the E1 enzyme in the ISGylation cascade. Through detailed NMR titration experiments and mutational studies, we mapped the interaction surface of Uba7-UFD with UbcH8, identifying key residues and their contributions to the E1-E2 interaction. We found that Uba7-UFD is flexible in its free form; this flexibility is conserved across ubiquitin-like systems but shows a regional shift that may play an important role in correct E2 and Ubl selection. Chemical shift perturbation and mutational analysis further demonstrate the importance of specific residues, particularly Cys996, in maintaining UFD’s structural integrity and binding capacity. Additionally, mutations designed to alter the flexibility and length of the loop region between UFD and UbcH8 show significant effects on binding, indicating that these regions are crucial for efficient E2 recruitment. These findings provide new insights into the mechanistic basis of E2 enzyme selection in ISGylation and underscore the functional relevance of dynamic structural transitions in E1-E2 complex formation.

**Statement of Significance:** ISGylation is a key ubiquitin-like modification pathway essential for antiviral defense, immune regulation, and protein quality control. However, the molecular principles that govern communication between the E1 and E2 enzymes in this pathway remain poorly defined. Here, we present the first NMR structure of the human Uba7 ubiquitin-fold domain (UFD) and its interaction with UbcH8. Our findings reveal how structural flexibility within the UFD is crucial for specific and efficient E2 recognition, providing fundamental insight into the dynamic mechanism of ISGylation and advancing our understanding of ubiquitin-like enzyme cascades relevant to cellular regulation and disease.

## Introduction

ISGylation is a post translational modification (PTM) that results in the binding of interferon-stimulated gene 15 (ISG15) to the target protein. ISG15 is a 15 kDa ubiquitin-like protein (Ubl) containing two ubiquitin-like domains connected by a short linker peptide (Haas et al., 1987). Expression of ISG15 is primarily induced by type I interferon (IFN α/β) signaling (Farrell et al., 1979), through interactions between IFN regulatory factors (IRFs) and interferon-stimulated response element (ISRE)-containing promoters (Sadler and Williams, 2018). Innate immune response on a molecular level is generated via host pattern-recognition receptors and expression of IFN-I through downstream signaling. In this regard, ISG15 expression coincides with initiation of the innate immune response which serves as the first line of defense against pathogens. After expression, ISG15 takes part in innate immune response either as an intracellular regulatory protein or through ISGylation of primarily pathogen proteins (Zhang et al., 2021). In the context of the innate immune response, effectiveness of ISGylation against pathogens varies between species. For instance, ISGylation in mice is shown to be a highly effective antiviral mechanism for inhibiting viral assembly slowing down pathogenesis (Durfee et al., 2010). ISGylation of viral proteins also interferes with localization, host-viral protein interactions, and protease activity thereby strengthening the antiviral response (Rahnefeld et al., 2014). Whereas in humans, inherited ISG15 deficiency only results in susceptibility to mycobacterial, not viral diseases indicating differing effectiveness against pathogens and possible differences in ISGylation pathway between species (Bogunovic et al., 2012).

Similar to ubiquitin, ISG15 is also covalently attached via its C-terminal LRLRGG motif to a lysine residue on the target protein through an analogous E1-E2-E3 enzymatic cascade (Narasimhan, Potter and Haas, 1996). ISGylation enzymatic cascade includes an activating E1enzyme (Uba7/Ube1L), a conjugating E2 enzyme (UbcH8), and an E3 ligase (HERC5) which their expression is simultaneously induced with ISG15 through ISRE containing elements. ISGylation begins with an ATP dependent reaction of sequential adenylation at the C-terminal motif “LRLRGG” generating an AMP-ISG15 intermediate which forms a thioester bond with the catalytic cysteine of the E1 activation enzyme, Uba7. In this step, active site cysteine half domains, the first catalytic cysteine half-domain (FCCH) and the second catalytic cysteine half-domain (SSCH) of the E1 form a thioester bond with the adenylated C-terminal glycine of ISG15. The ubiquitin folding domain (UFD) of Uba7 functions as the recognizing agent of the E2 enzyme and ISG15 where ISG15 is transferred to the catalytic cysteine of the E2 conjugating enzyme, UbcH8, through a transthioesterification reaction. E2 functions as a bridge between the Uba7 (E1), and HERC5 (E3), enzymes containing catalytic domain interaction sites of both E1 and E3. In the subsequent reaction, ISG15 is transferred to the catalytic cysteine of E3 through another transthioesterification reaction then ligated to a lysine residue of the target protein in a nonspecific manner (Haas, 2007). Despite its similarities, ISGylation differs considerably from ubiquitylation in terms of variability and prevalence in the cell. Unlike its counterpart, ISGylation cascade contains only one set of E1-E2 enzymes and therefore regulation of ISGylation is further influenced through intrinsic interactions between enzymes of the cascade in addition to external pathways regulating the ISGylation.

As the cornerstone of Ub/Ubl conjugation pathways, E1 enzymes are responsible for initiating the activation of the corresponding Ub/Ubl along with its transfer to the relevant E2 enzymes. This crucial function helps to ensure the accuracy and reliability of Ub/Ubl-mediated conjugation pathways (Capili and Lima 2007, Streich and Haas, 2010) highlighting the crucial role that E1 enzymes play in maintaining the integrity of ubiquitylation pathways. Previous studies havemanaged to identify crystal structures of Ub E1 along with E1s for Ub-like modifiers (Ubls) such as SUMO and Nedd8 (Figure 1), both singularly and in complex states with their corresponding Ubls (Olsen and Lima, 2013; Huang et al., 2005, Lois and Lima, 2005). Structures of the SUMO E1 provide mechanistic insights into SUMO activation and E2 recruitment to E1. A discovery from these studies was the revealing of the multidomain structure inherent to canonical E1s. The domains include a pseudodimeric adenylation domain which plays a crucial role in Ubl activation, a Cys domain which contains the catalytic cysteine residue and a ubiquitin-fold domain (UFD) involved in the recruitment of E2 (Figure 1). To better contextualize the structural findings of the present study, we first conducted a comparative analysis of E1 enzyme ubiquitin-fold domains (UFDs) from several ubiquitin and ubiquitin-like (Ubl) modification systems (Figure 1). The aim of this figure is to illustrate that in most available crystal structures of free (apo) E1 enzymes, the E2-binding regions of the UFD domains lack interpretable electron density, indicating substantial conformational flexibility in these segments. This absence of density across multiple E1–UFD structures—including Uba1, Uba2, Uba3, and Uba6—suggests that these regions are inherently disordered or mobile in solution until they engage their cognate E2 enzymes. Such disorder-to-order transitions have been previously proposed to facilitate accurate E2 selection and dynamic repositioning of the catalytic domains during transthiolation. Our NMR-derived structure of the human Uba7-UFD (Figure 1K) provides complementary insight into this phenomenon by capturing the domain in its solution state, where flexibility is preserved rather than averaged out as in crystal lattices. In constructing Figure 1, we intentionally combined representative E1–UFD structures, both apo and complexed with E2 or Ubl partners, to highlight how the flexible β-grasp and loop regions adopt defined conformations upon complex formation.

**Figure 1.**
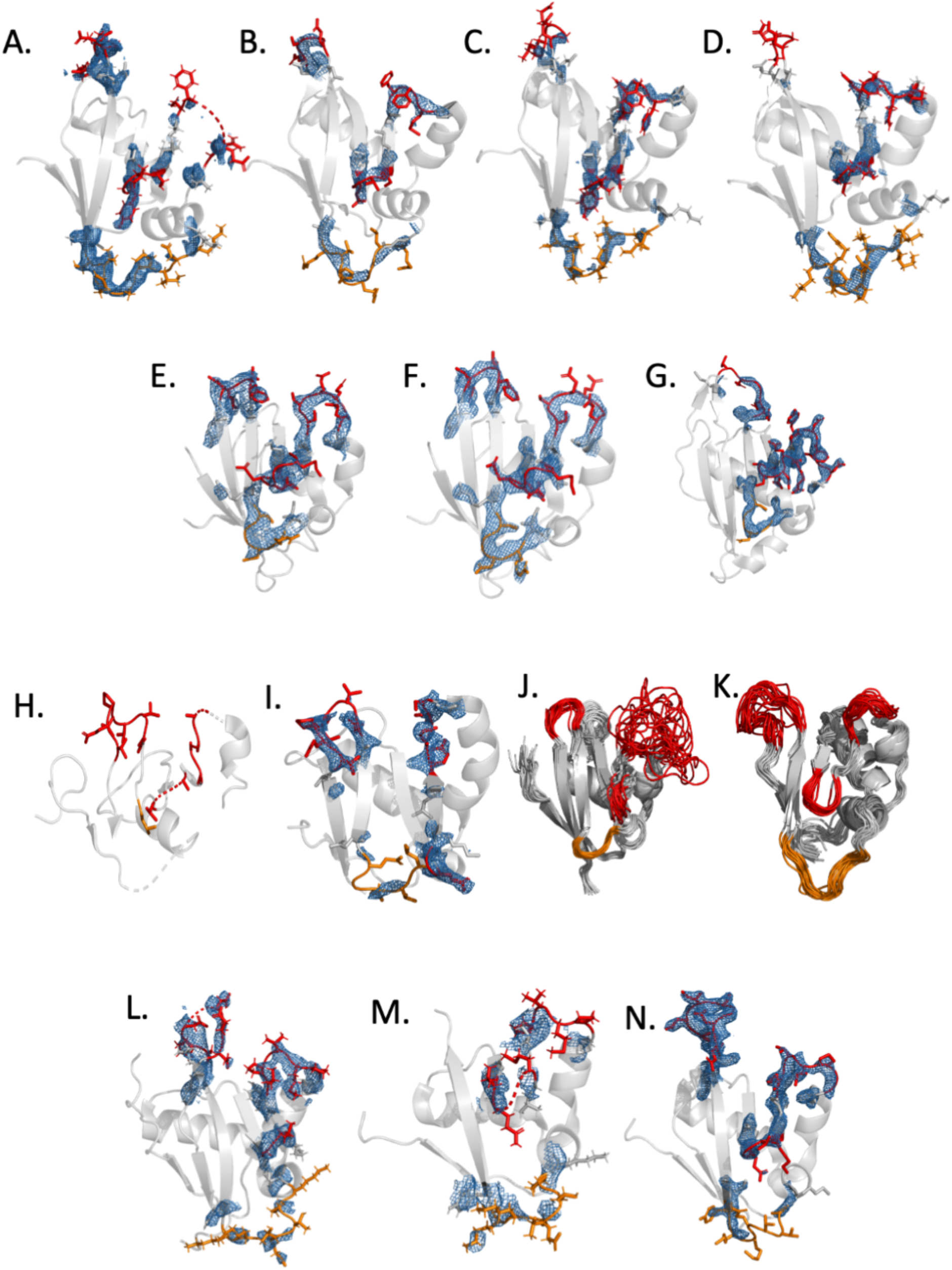
Ubiquitin and Ubiquitin-like post-translational modification systems E1 enzymes’ ubiquitin fold domain structures. Regions on the structures that have high plasticity and directly interact with E2 are marked in red, and regions thought to interact with Ubl are marked in orange. Electron density maps are also shown over the colored regions. A. Saccharomyces cerevisiae Uba1 Apo form (PDB: 7ZH9) B. Homo sapiens Uba1 complex with ubiquitin (PDB: 6DC6) C. Saccharomyces cerevisiae complex with Ubc13 (E2) (PDB: 6ZHS) D. Schizosaccharomyces pombe complex with Ubc15 (E2) and ubiquitin (PDB: 5KNL) E. Homo sapiens Uba2 apo form (6CWZ) F. Homo sapiens Uba2 complex with SUMO (3KYC) G. Saccharomyces cerevisiae Uba2-UFD complex with Ubc9 (3ONG) H. Homo sapiens Uba3 apo form (1YOV) I. Homo sapiens Uba3-UFD complex with Ubc12 (1Y8X) J. Homo sapiens Uba3 UFD apo form (2LQ7) K. Homo sapiens Uba7-UFD apo form (8WWX) L. Homo sapiens apo Uba6 (7PVN) M. Homo sapiens Uba6 complex with FAT10 (7PYV) N. Homo sapiens Uba6 complex with FAT10 (7SOL)

Furthermore, these prior structural studies have unveiled the considerable conformational shifts E1s undergo to execute their roles effectively. In light of conformational changes revealing the UFD, a hidden E2 binding area on the E1 is exposed, assisting in facilitating contact between the Ubl thioester and E2 (Olsen and Lima, 2013). Upon the Ubl thioester’s transfer from E1 to E2, it is proposed that the UFD reverts to its originally locked conformation. This switch aids in the release of the E2-Ub thioester product through a spatial mechanism (Olsen and Lima, 2013). Similarly, E1-E2 interaction during ISGylation marks a crucial regulatory step of the ISGylation pathway however while considerable insights have been attained regarding the structural insights into E1 activity, the specific process at the atomic level has yet to be determined of ISGylation pathway. The importance of flexible regions and flexibility in protein-protein interactions has become increasingly evident with the advancement of structural methods. In this context, “disorder-to-order transition” in protein binding regions is being extensively studied using NMR spectroscopy and plays a crucial role in elucidating protein-protein interactions (Jubb, Blundell and Ascher, 2015). Specific mechanistic and interactive characteristics of ISGylation are still poorly understood. In this study, we explore the NMR structure of Uba7-UFD, interaction dynamics and the critical role of the disorder-to-order transition within the E1-E2 complex of the ISGylation cascade at the residual level using NMR spectroscopy. The structural flexibility observed for Uba7-UFD in the unbound state is consistent with a general feature of E1 enzymes, in which localized dynamic elements are maintained within an otherwise well-folded UFD scaffold to support productive E2 engagement. Upon interaction with UbcH8, the relaxation profiles indicate a redistribution of backbone dynamics within the UFD—most prominently around the highlighted β2/β4 surface—suggesting that E2 binding modulates pre-existing motional properties rather than invoking a global conformational switch. Notably, while flexibility at the UFD–E2 interface appears conserved across ubiquitin-like activation systems, the specific segments exhibiting the strongest dynamic signatures show a measurable regional shift between different E1–E2 pairs, providing a plausible mechanism for fine-tuning E2 specificity and ensuring efficient ISG15 transfer. Through detailed NMR titration experiments, we mapped the extensive interaction surfaces between Uba7-UFD and UbcH8, identifying over 30 residues involved in this dynamic binding process. The study also highlighted the significance of electrostatic complementarities between the interacting regions of Uba7-UFD and UbcH8, which contribute to their stable complex formation. Furthermore, mutational analysis underscored the importance of key residues, such as cysteine 996, in maintaining the structural integrity and binding capacity of Uba7-UFD. These findings provide novel insights into the conformational plasticity of E1 enzymes and their regulatory role in ISGylation, paving the way for a deeper understanding of this post-translational modification process at the molecular level.

## Materials and Methods

### Construction of Uba7-UFD expression plasmid

The DNA sequence coding for Uba7-UFD (amino acid residues 903-1011) was amplified from pGEX-full length HsUba7 by polymerase chain reaction (PCR). Taq polymerase (Thermo) and primers 5’ Uba7-UFD-BglII (5’-GCGCAGATCTCAGACGTTCCATCACCTG-3’) and 3’ Uba7-UFD-HindIII 5’-GCGCAAGCTTACAGCTCATAGTGCAGAGG-3’ were used in the reaction. Amplified PCR product was cleaned up using Omega PCR clean-up kit and digested with BglII and HindIII while the pQE30-GBI expression vector was digested with BamHI and HindIII. The digested PCR product and plasmid ligated with T4 DNA ligase. Accuracy of the cloned gene was confirmed by DNA sequencing (Macrogen, Korea) using reverse and forward pQE30 primers. Recombinant pQE-30-GBI-Uba7-UFD plasmid transformed into the chemically competent SG13009 expression strain of *E. coli* using heat shock.

### Site-Directed mutagenesis to Uba7-UFD

For site-directed mutagenesis PCR reaction, Pfu polymerase was mixed with template pQE-GBI-Ube1-UFD plasmid and forward and reverse primers containing the desired mutation. Primer names and sequences used for mutations are given below.

*5’ C996A 5’ GTGCTAGAGCTGAGCGCTGAGGGTGACGACG 3’*

*3’ C996A 5’ CGTCGTCACCCTCAGCGCTCAGCTCTAGCAC 3’*

*5’ C996S 5’ GTGCTAGAGCTGAGCAGCGAGGGTGACGACG 3’*

*3’ C996S 5’ CGTCGTCACCCTCTCGGCTCAGCTCTAGCAC 3’*

*5’ A1004C 5’ GGGTGACGACGAGGACACTTGCTTCCCACCTCTGCACTATG 3’*

*3’ A1004C 5’ CATAGTGCAGAGGTGGGAAGCAAGTGTCCTCGTCGTCACCC 3’*

Once completed, PCR reaction mixture was incubated with 10 U DpnI restriction enzyme for 2 hours at 37 °C to digest the template plasmid. Resulting reaction mixture containing amplified mutant plasmid was transformed into Novablue competent *E. coli* cells. The existence of the desired mutation was confirmed by DNA sequencing. Plasmids containing target mutation were selected and transformed into competent SG13009 (pREP4) *E. coli* cells. Plasmids encoding mutant R944L and 998GGG proteins were purchased from GENSCRIPT.

### Expression of recombinant GBI-Uba7-UFD fusion protein in *E. Coli*

The freshly-streaked single colony of SG13009 strain of *E. coli* containing PQE30-GBI-Uba7-UFD plasmid was inoculated in 10 mL Luria Broth (LB) media containing 150 μg/mL ampicillin and 50 μg/mL kanamycin. Inoculated culture was grown overnight with shaking at 250 rpm at 37°C. Next morning, 10 mL overnight culture was added to 1 L fresh LB media containing the same concentrations of antibiotics and grown at 37 °C and 250 rpm until OD_600_ reached 0.8. The protein expression was induced with 1 mM IPTG. After induction, cells were grown for 5 more hours at 37 °C. Cells were then centrifuged at 4500 x g for 25 minutes at 4 °C and stored at – 80 °C until protein purification.

### Refolding and purification of fusion protein with affinity chromatography

For purification of GBI-Uba7-UFD fusion protein, cell paste was resuspended in 30 mL Lysis buffer containing 50 mM Tris (pH 7.5), 150 mM NaCl, 5 mM DTT and sonicated at 4 °C, then centrifuged at 15000 x g for 60 minutes at 4 °C. Proteins in pellet and supernatant were checked by SDS-PAGE and supernatant were discarded. Inclusion bodies in pellet were washed two times with 50 mL 0.5% Triton X-100 in 50 mM Tris (pH 7.5), 150 mM NaCl to solubilize membranes and membrane proteins and then centrifuged at 15000 x g for 60 minutes at 4 °C. Cleaned inclusion bodies were solubilized in 100 mL denaturation buffer containing 7 M Guanidium-HCl, 50 mM Tris, 150 mM NaCl, 10 mM DTT overnight with stirring at 20 °C. The next day, the mixture was centrifuged at 15000 x g for 60 minutes at 15 °C to separate insoluble components from the solution. Refolding of fusion protein was achieved by dialysis. For this, supernatant was transferred into a dialysis membrane (MWCO value 3500) and dialyzed twice against 4 liters of 50 mM Tris, 150 mM NaCl, 10 mM DTT pH 7.5 buffer for 8 hours at 15 °C. After second dialysis, precipitated components were removed by centrifugation at 15000 x g for 60 minutes at 15 °C. Refolded fusion protein was purified by immobilized metal ion affinity (IMAC) chromatography using a 5 mL HisTrap HP column (GE HealthSciences) precharged with nickel and attached on an Akta Prime Plus protein purification system. The dialyzed sample was bound to a HisTrap HP column at a flow rate of 3 mL/min. Unbound proteins were washed away with a buffer containing 50 mM Tris-HCl, 150 mM NaCl, 5 mM DTT, 20 mM imidazole pH 7.5 until the absorbance at 280 nm was below 10 mAbs. Elution of HisTagged fusion protein was achieved by linearly increasing the amount of elution buffer (50 mM Tris, 150 mM NaCl, 5 mM DTT, 500 mM Imidazole pH 7.5) from 0 to 100 % in 30 mL volume. All fractions were analyzed by SDS-PAGE and those containing the purified GBI-Uba7-UFD fusion protein were pooled, concentrated and desalted with 20 mM phosphate Buffer pH: 7.4 to 200 μM volume in a pressure concentrator (Stirred cell, Amicon) using a membrane with 3500 Da MWCO.

### Isotopically labeled protein expression

For 1 L of M9 minimal media 200 mL sterile 5x M9 salts lacking ^15^N-NH_4_Cl (64 g Na_2_HPO_4_ 7H_2_O, 15 g KH_2_PO_4_, 2.5 g NaCl), were diluted to 950 mL with dd H_2_O. 1 g ^15^N-NH_4_Cl (dissolved in 10 mL ddH_2_O and filter sterilized), 2 mL sterile 1 M MgSO_4_, 20 mL 20% filter sterilized glucose (for double labeled protein samples 2 g/L ^13^C glucose), and 0.1 mL sterile 1M CaCl_2_ was added to M9 salts and the volume was completed to 1 L with sterile ddH_2_O. Media was then supplemented with trace metals and vitamins.

Freshly transformed cells were used to inoculate 20 mL of LB media containing ampicillin and kanamycin. Cultures were grown overnight at 37 °C, 250 rpm. Cells were pelleted the next day by centrifugation at 1000 x g for 10 min at 4 °C, resuspended in 10 mL of M9 minimal media and added to the 1L of either ^15^N-labeled M9 culture for ^15^N-labeled protein production or ^15^N-^13^C labeled M9 culture for double-labeled protein production. Protein expression was carried out as described above with the exception that after induction with 0.5 mM IPTG, cells were incubated at 25 °C for 15 hours to allow for expression of target proteins. Cells were then harvested by centrifugation at 4000 x g for 30 minutes. Cell paste was kept at -20 °C until further processing.

### NMR data collection

2D ^1^H-^15^N HSQC, 3D HNCA, 3D SE HNCO, 3D SE HNCACB, 3D SE HN(CO)CA, 3D SE CCONH, 3D SE HCCONH, 3D SE HBHACONH, 3D HNCACO, 3D SE HNCOCACB, 3D HCCH-TOCSY 3D SE ^15^N-NOESY-HSQC, 3D SE ^13^C-NOESY-HSQC-aliphatics and 3D ^13^C-NOESY-HSQC-aromatics were performed at 20 ^0^C on Bruker Avance II 600 MHz spectrometer with a triple resonance z-axis gradient CryoProbe. All NMR samples were prepared in 20 mM sodium phosphate, pH 7.4 containing 5-10% D_2_O and 0.05% sodium azide.

### NMR data processing, analysis and structure calculation

All multidimensional NMR datasets were processed using NMRPipe (Delaglio et al., 1995). Raw free-induction decay (FID) data were subjected to zero-filling to at least twice the acquired complex points, apodized with a shifted sine-bell window, and Fourier-transformed in both dimensions. Phase and baseline corrections were applied manually. Processed spectra were visualized and analyzed using XEASY and the POKY suite (Lee et al., 2021). Backbone and side-chain resonance assignments were obtained from standard 3D triple-resonance experiments (HNCA, HNCO, HNCACB, HN(CO)CA, CCONH, HCCONH, HBHACONH, HCCH-TOCSY,

15N-NOESY-HSQC, and 13C-NOESY-HSQC). Secondary-structure elements were identified from chemical-shift indices using CSI and torsion-angle restraints derived with TALOS+ (Shen et al., 2009). Inter-residue NOE distance restraints were extracted from NOESY spectra and combined with dihedral-angle restraints for structure calculations in POKY/CNS. The final ensemble of 20 lowest-energy conformers was refined against all experimental restraints and validated using standard NMR structure-quality metrics (RMSD, Ramachandran, and NOE statistics).

All NMR titration experiments and HSQC spectra were acquired using isotopically labeled protein samples at a concentration of 200 μM. HSQC cross-peak movements were quantified as a combined ^1^H-^15^N chemical shift perturbation calculated from ΔδAV = [(Δδ ^1^H*5)^2^ + (Δδ ^15^N)^2^]^½^ equation. The structure calculations were performed using POKY (Lee et al., 2021) programs on a computer running the MacOS operating system.

### Docking Analysis

The docking process in HADDOCK2.4 docking protocol (de Vries et al., 2010) is divided into three stages. In the first stage, the proteins are treated as rigid bodies, and their interaction energy is calculated for different orientations. The orientations with the lowest energy are then selected for further refinement. In the second stage, simulated annealing is used to refine the interface between the two proteins. Simulated annealing is a Monte Carlo method that allows the system to escape from local energy minima. In the context of protein-protein docking, simulated annealing allows the proteins to explore different conformations at the interface and identify the conformation with the most favorable energy. In the final stage, the complex is immersed in solvent to improve the energetics. The solvent molecules are explicitly included in the simulation, and their interactions with the proteins are taken into account. This stage helps to refine the structure of the complex and ensure that it is stable in a solvated environment. By combining these three stages, HADDOCK2.4 outputs a prediction of the structure of protein-protein complex.

### NMR relaxation measurements

Backbone dynamics of Uba7-UFD and UbcH8 were characterized using ^15^N relaxation experiments (T_1_, T_2_, and {^1^H}-^15^N heteronuclear NOE) to assess conformational flexibility and dynamic changes upon complex formation. Relaxation measurements were performed for the free ^15^N-labeled Uba7-UFD and free ^15^N-labeled UbcH8 under identical buffer conditions (20 mM phosphate buffer, pH 7.4, 150 mM NaCl, 5 mM DTT, 5–10% D_2_O).

Experiments for Uba7-UFD were carried out at 20 °C on a Bruker Avance III 900 MHz spectrometer equipped with a triple-resonance cryogenic probe, while UbcH8 relaxation data were acquired under identical conditions on a Bruker Avance II 600 MHz spectrometer.

Standard Bruker pulse sequences were used for all relaxation measurements. T_1_ relaxation times were obtained using eight relaxation delays (10, 50, 100, 200, 400, 600, 800, and 1000 ms), and T_2_ relaxation times were recorded using eight delays (10, 30, 50, 70, 90, 110, 130, and 150 ms). {^1^H}-^15^N heteronuclear NOE spectra were collected with and without proton presaturation (3 s) using identical acquisition parameters. All spectra were processed with NMRPipe and analyzed using Bruker Dynamics Center software.

## Results

### Backbone and side-chain resonance assignments of native human Uba7-UFD

Uba7-UFD residues 903-1011 were recombinantly produced with an N-terminal GBI fusion in *E. coli* cells and purified using Ni^2+^ affinity chromatography. Despite efforts to produce a soluble construct, the GBI-Uba7-UFD expressed into inclusion bodies similar to our previous study with Uba3-UFD (Elgin et al., 2012). The most effective approach to refold GBI-Uba7-UFD was denaturation with 7 M guanidinium chloride and 5 mM DTT followed by dialysis into a solution containing 50 mM Tris, 150 mM NaCl, 5 mM DTT, pH 7.5. Unfortunately, Uba7-UFD possessed very low stability/solubility once the GBI solubility tag was removed resulting in precipitation within ∼4 h (data not shown). We next evaluated the possibility of using GBI-Uba7-UFD fusion for NMR structure and binding studies. Comparison of 2D ^1^H-^15^N HSQC spectra for both the free domain and the GBI fusion showed no detectable difference in the amide resonances demonstrating the fusion does not affect the Uba7-UFD tertiary structure (Supp Fig. 1). Comparison of ^1^H-^15^N HSQC spectra collected before (day 1) and after (day 9) a suite of 3D assignment experiments reveals no significant time-dependent changes (Supp Fig. 2). Therefore, GBI-Uba7-UFD was found to be suitable for further structure data collection and titration experiments.

**Figure 2.**
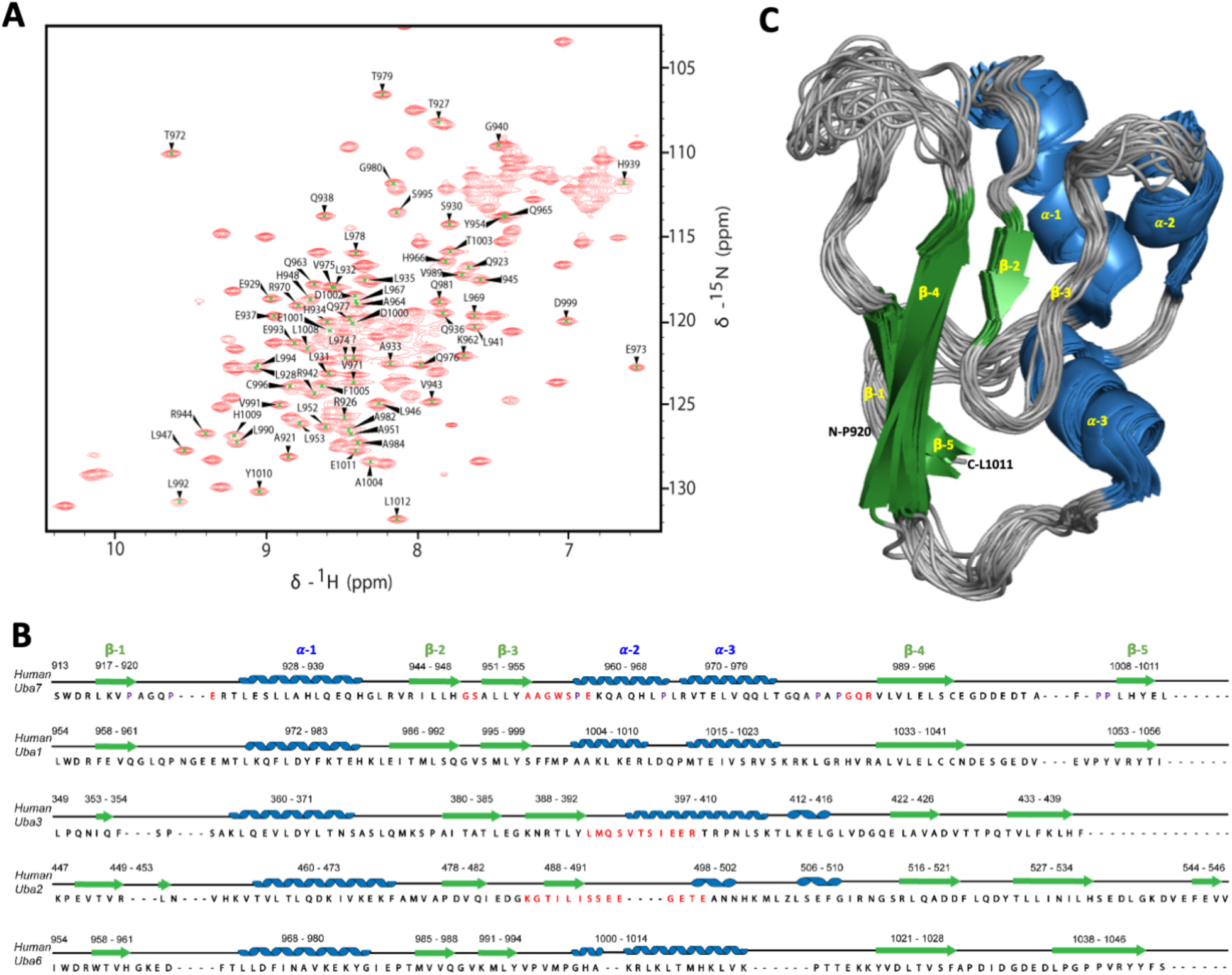
NMR structure of Human Uba7 ubiquitin fold domain A. Human Uba7 enzyme ubiquitin fold domain ^1^H-^15^N HSQC spectrum and N-H assignments. B. Human Ubiquitin-like systems activating enzymes ubiquitin fold domain sequential and secondary structure alignment C. Superposition of the 20 lowest-energy NMR structures of the Uba7 ubiquitin fold domain

The chemical shift dispersion of [*U*-^15^N]-GBI-Uba7-UFD amide resonances are consistent with a well-folded protein (Fig. 2A); however, only 71 of the 100 expected backbone N-H resonances were observed (109 total residues including eight prolines). These were assigned to residues 919-1011 using a suite of conventional 3D triple-resonance spectra from which secondary structure was predicted using TALOS+ (Shen et al., 2009) (Fig. 2A). A stretch of seven residues following the third beta-strand were unassignable. The homologous region of the two E1 enzyme UFDs, Uba3 and Uba2 (Wang et al., 2009; Elgin et al., 2012), were similarly unassigned due to low signal intensity in 3D experiments which suggests these missing peaks result from inherent plasticity of these regions (Fig. 2B).

### NMR Structure determination of Human Uba7-UFD

The three-dimensional structure of the human Uba7-UFD was successfully determined using multi-dimensional NMR spectroscopy (Fig. 2C). A total of 1616 unique NOE distance and 153 (phi/psi) dihedral angle constraints are used in the final structure calculations; refinement statistics for the ensemble of 20 lowest energy conformers are shown in Table 1. The final ensemble exhibited good convergence with an average pairwise root mean square deviation (RMSD) of 0.9 Å for backbone atoms and 1.4 Å for all heavy atoms. The secondary structure consists of three α-helices and four β-strands arranged in a ubiquitin fold domain (Fig. 2B, 2C; Supp Fig. 4) consistent with previously reported ubiquitin-like domains (Wang et al., 2010; Lv et at., 2018). After the second β-strand, a partially folded intermediate third β-strand is present, with distinct loops located before and after it. These loops and β-strand may play a role in the interaction with the E2 enzyme in the ISGylation cascade, similar to the mechanism observed in ubiquitination (Lv et al., 2018). The crystal structures of apo E1 enzymes (PDB:7ZH9, 6CWZ, 7PVB) lack electron density in certain portions of the UFDs (Walden, Podgorski, and Schulman, 2003) (Figure 1). This is consistent with NMR analyses that the UFDs contain intrinsically disordered regions in their free form (Wang et al., 2009; Elgin et al., 2012) that adopt stable secondary structures upon E1-E2 complexation (Huang et al., 2005; Wang et al., 2010) (Figure 1). Comparison of NMR structures of UFD’s from Uba7 and Uba3 shows that Uba7-UFD NMR structure shows a well-folded second alpha helix (Figure 2B-Uba7_960-968), which is largely unfolded in Uba3-UFD (Figure 2B-Uba3_397-403) (PDB 2LQ7; Fig. 1J-K,2B). In contrast, the third beta strand of Uba7-UFD (Figure 2B-Uba7_950-960), which is well defined in Uba3-UFD (Figure 2B-Uba3_388-392), is relatively flexible (Fig. 1H-K). In addition, the NMR-derived structure showed that the acidic loop region located between the fourth and fifth beta strands also exhibits much greater flexibility compared to the corresponding loop region in Uba3-UFD (Fig. 1H-K). Although the overall ubiquitin fold structure is conserved in both Uba7- and Uba3-UFDs, NMR ensembles clearly demonstrate that the location of conformational flexibility is quite unique to each domain.

**Table 1.** Restrains and statistics for 20 best solution NMR structures of Uba7-UFD.

|  |  |
| --- | --- |
| **Distance and dihedral restraints** |  |
| Short range ($\vert i-j\vert \leq 1$) | 224 |
| Medium range ($\vert i-j\vert < 5$) | 319 |
| Long range ($\vert i-j\vert > 5$) | 358 |
| Dihedral restraints ($\phi/\psi$) | |
| Dihedral restrains | 153 |
| Total number of restricting constraints | 1616 |
| **Average RMSD (Å) against the lowest energy model for ordered residues (PSVS)** |  |
| Backbone | 0.9 |
| All heavy atoms | 1.4 |
| **RMSD from ideal geometry** |  |
| Bond lengths (Å) | 0.015 |
| Bond angles (°) | 1.4 |
| **Consistent violations** |  |
| Distance constraints ($>0.5$ Å) | 0 |
| Dihedral angle constraints ($>5^\circ$) | 14.6 |
| Van der Waals constraints ($>0.2$ Å) | 1.8 |
| **Ramachandran analysis** |  |
| Favoured regions (%) | 93.7 |
| Allowed regions (%) | 6.2 |
| Generously allowed | 0.1 |
| Disallowed | 0 |

### Chemical shift mapping reveals the Uba7-UbcH8 interface

To test the hypothesis that E1-UFD conformational disorder plays a role in E2 binding/selection/specificity, we performed an NMR titration to map UbcH8 onto the Uba7-UFD surface. ^1^H-^15^N HSQC spectra were collected on [*U*-^15^N]-Uba7-UFD mixed with increasing concentrations of unlabeled-UbcH8 in molar ratios of 1:0.25, 1:0.5, 1:1 and 1:1.75 (Supp Fig. 5). Inspection of these spectra revealed a subset of peaks that were dose-dependently perturbed upon addition of UbcH8 (Figure 3A); combined ^1^H/^15^N chemical shift changes were calculated and the most perturbed residues were highlighted onto the Uba7-UFD structure (Figure 3B, 3C). UbcH8 binding surface of Uba7-UFD appears to be quite large, involving about 18 residues. Interestingly, some of the peaks displaying the largest chemical shift perturbations upon addition of UbcH8 were weaker in intensity compared to the average peak intensities in the spectrum of the free Uba7-UFD. In fact, four peaks with pronounced chemical shift perturbations corresponded to those residues in the unassigned fragments and were absent in 3D spectra. One of the simplest explanations for the low intensity and absence in 3D spectra is that these residues may undergo conformational exchange on the microsecond-millisecond timescale (Supp Fig. 6).

**Figure 3.**
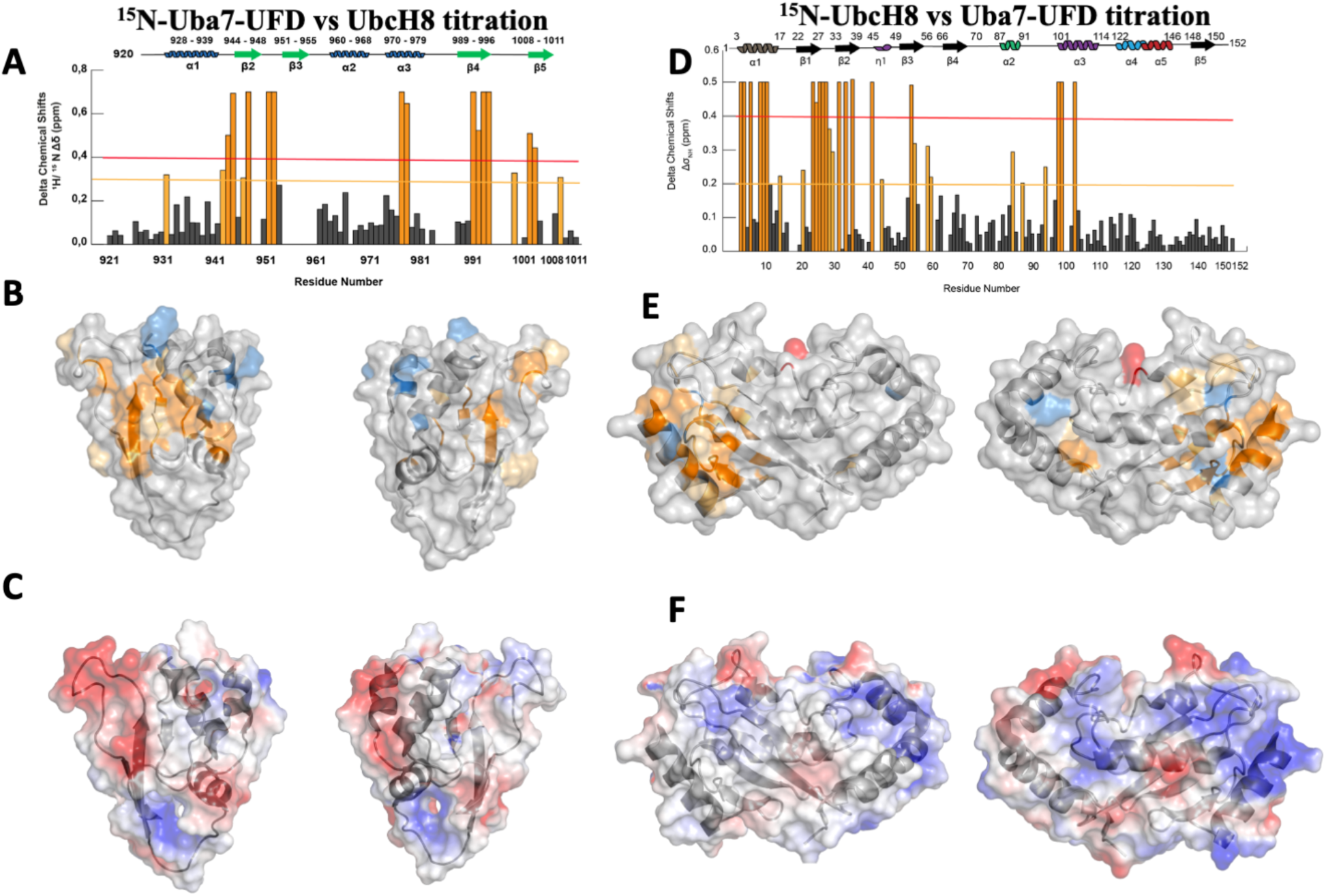
Mapping of interactions between UbcH8 and Uba7-UFD at the amino acid level and NMR data driven modeling of the Uba7-UbcH8 complex structure A. ^15^N-Uba7-UFD vs UbcH8 titration Chemical Shift Perturbation graph ^1^H-^15^N chemical shift perturbations for Uba7-UFD were calculated for each residue. The residues colored orange possessed chemical shift perturbations larger than 0.4 ppm threshold (red line). The residue colored merigold possessed chemical shift perturbations larger than the 0.3 ppm threshold (merigold line). B. ^15^N-Uba7-UFD vs UbcH8 titration Chemical Shift Perturbation map Residues with chemical shift perturbations larger than the 0.4 ppm threshold (red), residues with chemical shift perturbations larger than 0.3 ppm threshold (merigold) and peak intensity changes that were larger than the threshold (blue) were mapped onto the Uba7-UFD structure C. Surface electrostatic potential of Uba7-UFD Red and blue coloured regions denote negative and positive charges, respectively. D. ^15^N-UbcH8 vs Uba7-UFD Titration Chemical Shift Perturbation graph ^1^H-^15^N chemical shift perturbations for UbcH8 were calculated for each residue. The residues colored orange possessed chemical shift perturbations larger than 0.4 ppm threshold (red line). The residues colored merigold possessed chemical shift perturbations larger than the 0.2 ppm threshold (merigold line). E. ^15^N-UbcH8 vs Uba7-UFD Titration Chemical Shift Perturbation map Residues with chemical shift perturbations larger than the 0.4 ppm threshold (orange), residues with chemical shift perturbations larger than 0.2 ppm threshold (merigold) and peak intensity changes that were larger than the threshold (blue) were mapped onto the UbcH8 crystal structure (PDB 1WZV), red residue represents the active-site cysteine (Cys85) of UbcH8 involved in transthiolation. F. Surface electrostatic potential of UbcH8 Red and blue coloured regions denote negative and positive charges, respectively.

**Figure 4.**
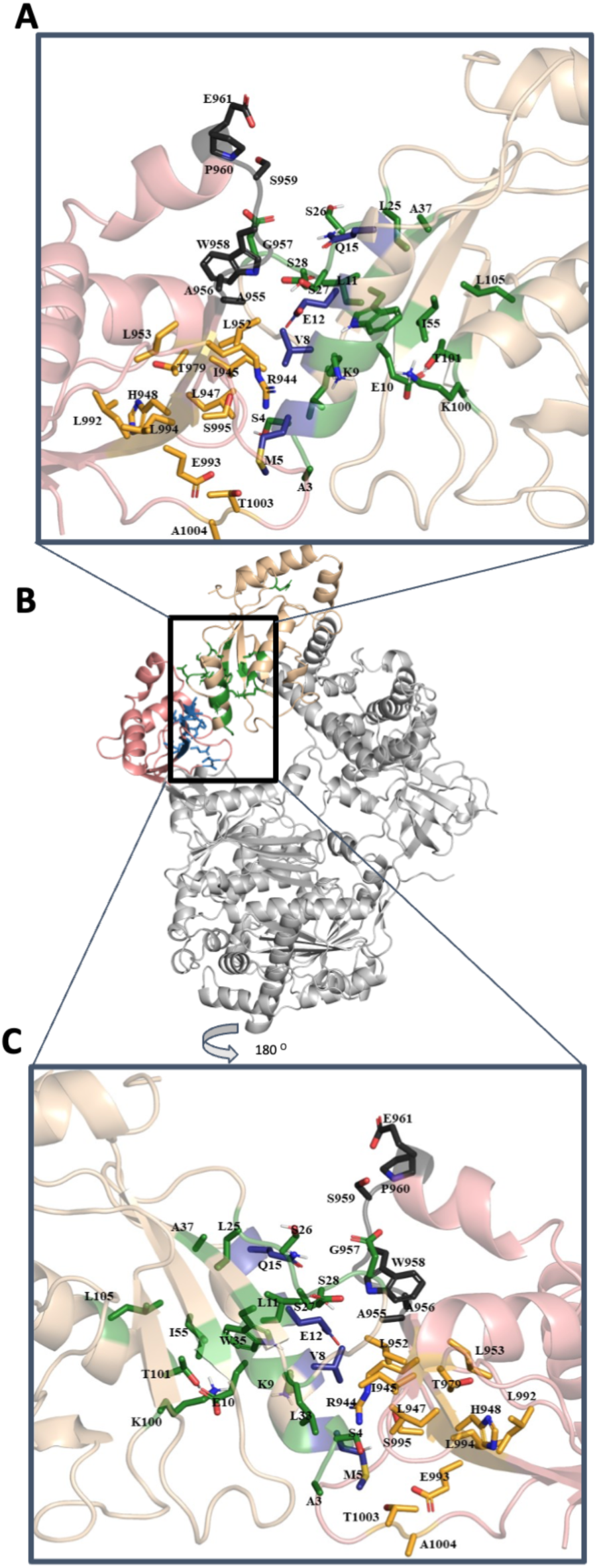
NMR data driven model of the Uba7-UbcH8 complex structure

**Figure 5.**
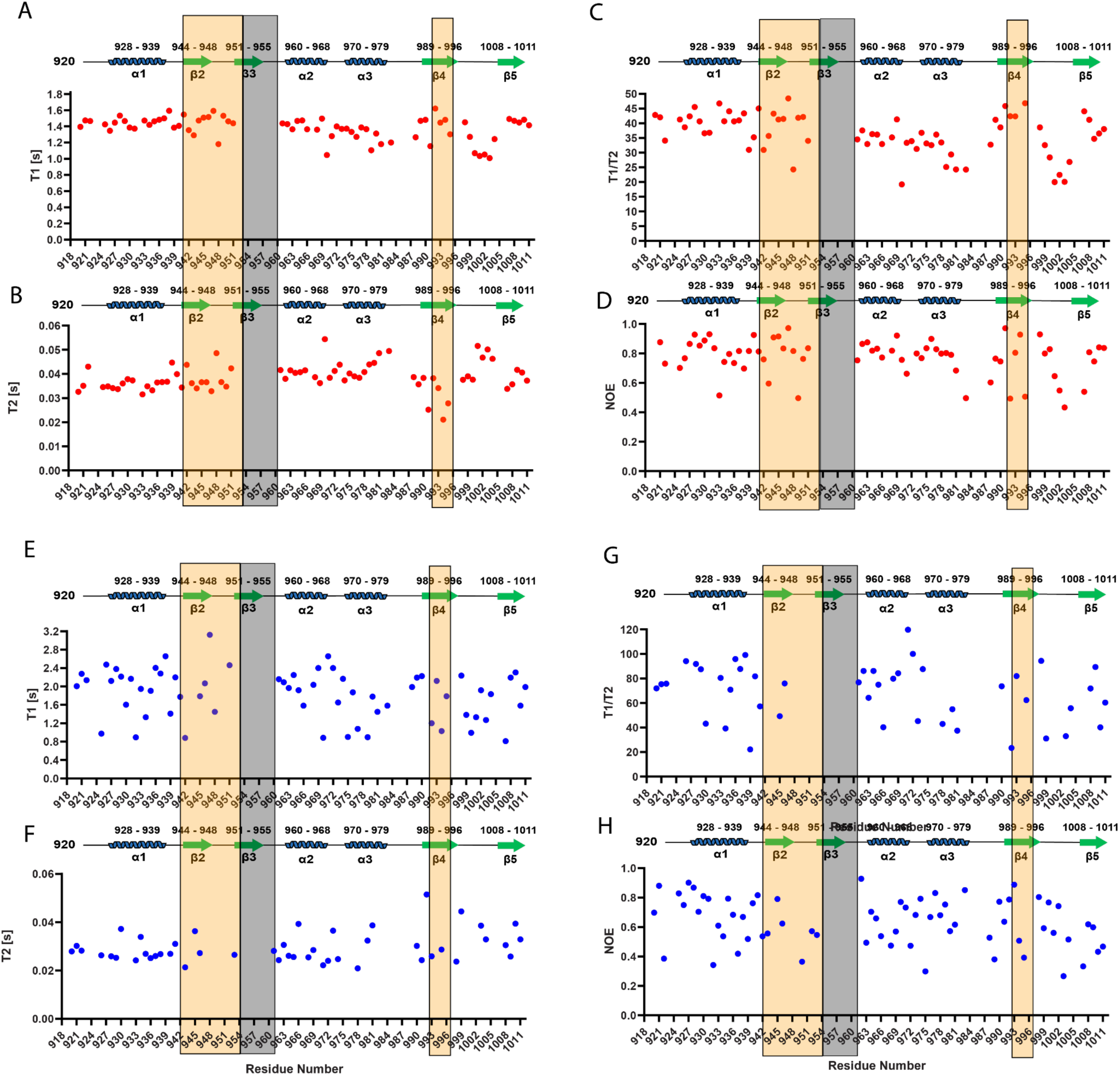
^15^N Relaxation analysis for Uba7 UFD and Uba7 UFD-UbcH8 complex A. ^15^N T1 relaxation rate as function of Uba7 UFD sequence B. ^15^N T2 relaxation rate as function of Uba7 UFD sequence C. T1/T2 values of Uba7 UFD protein as function of Uba7 UFD sequence D. ^1^H-^15^N NOE relaxation analysis for Uba7 UFD E. ^15^N T1 relaxation rates of ^15^N-labeled Uba7 in complex with unlabeled UbcH8 F. ^15^N T2 relaxation rates of ^15^N-labeled Uba7 in complex with unlabeled UbcH8 G. T1/T2 relaxation values of ^15^N-labeled Uba7 UFD in complex with unlabeled UbcH8, plotted as a function of the Uba7 UFD sequence H. ^1^H-^15^N NOE relaxation analysis for Uba7 UFD in complex with unlabeled UbcH8

**Figure 6.**
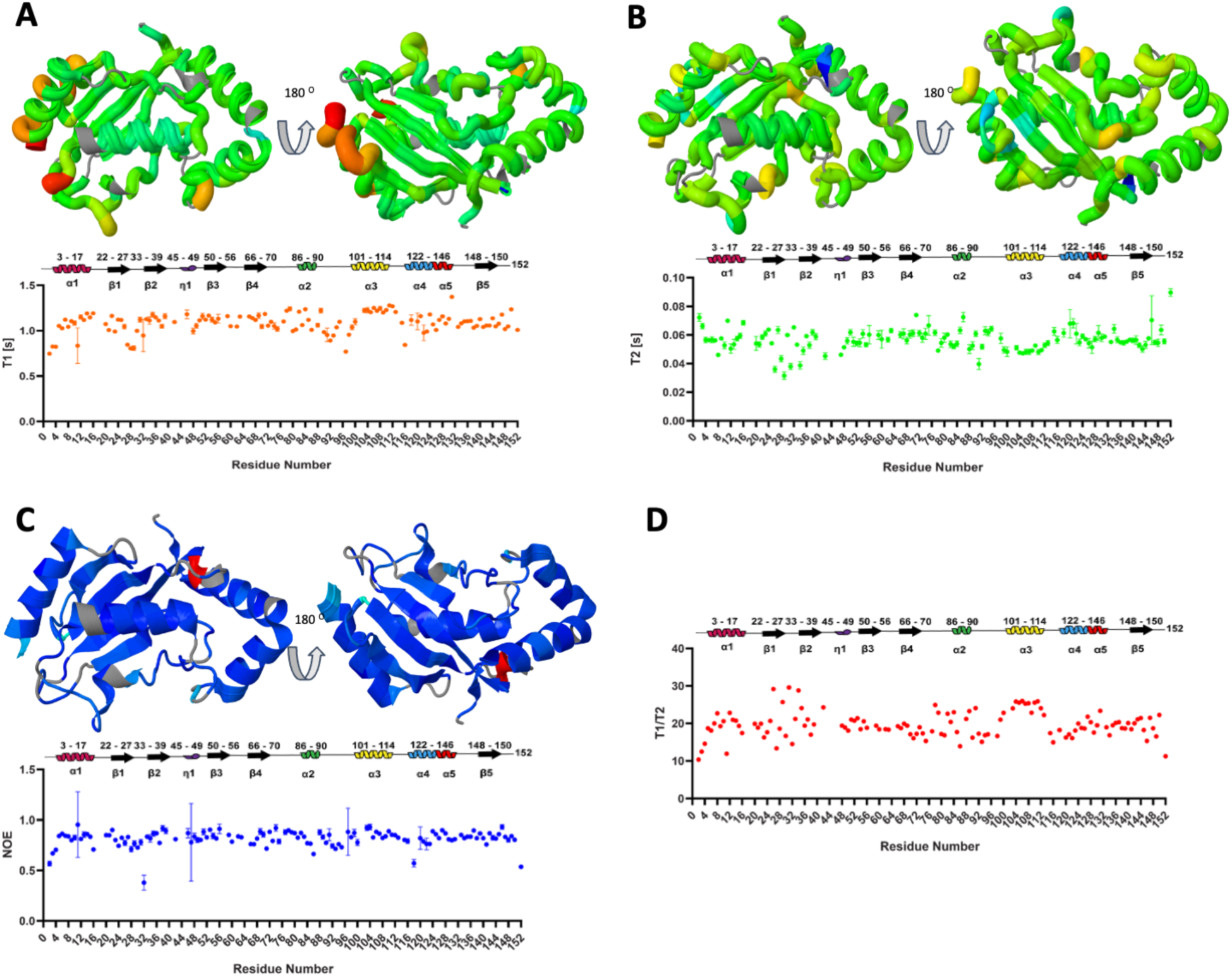
^15^N Relaxation analysis for UbcH8 A. ^15^N T1 relaxation rate as function of protein sequence and mapped on crystal structure (PDB:1WZW) B. ^15^N T2 relaxation rate as function of protein sequence and mapped on crystal structure (PDB:1WZW) C. ^1^H-^15^N NOE relaxation analysis for UbcH8 and mapped on crystal structure (PDB:1WZW) D. T1/T2 values of UbcH8 protein as function of protein sequence

We next mapped the binding interface onto the UbcH8 surface by titrating increasing concentrations of Uba7-UFD into [*U*-^15^N]-UbcH8. Substantial chemical shift perturbations clustered to the UbcH8 N-terminal region, helix 1, strands 1-3, and helix 3 (Figure 3D-F). Several peaks were also noted to disappear in the N-terminal region, helix 1, strands 1-3, and helix 3 upon addition of unlabeled Uba7-UFD (Supp Fig. 7). This result is consistent with the mutational study showing that the N-terminal region of UbcH8 is important in Uba7 interaction (Huibregtse et al. 2008).

**Figure 7.**
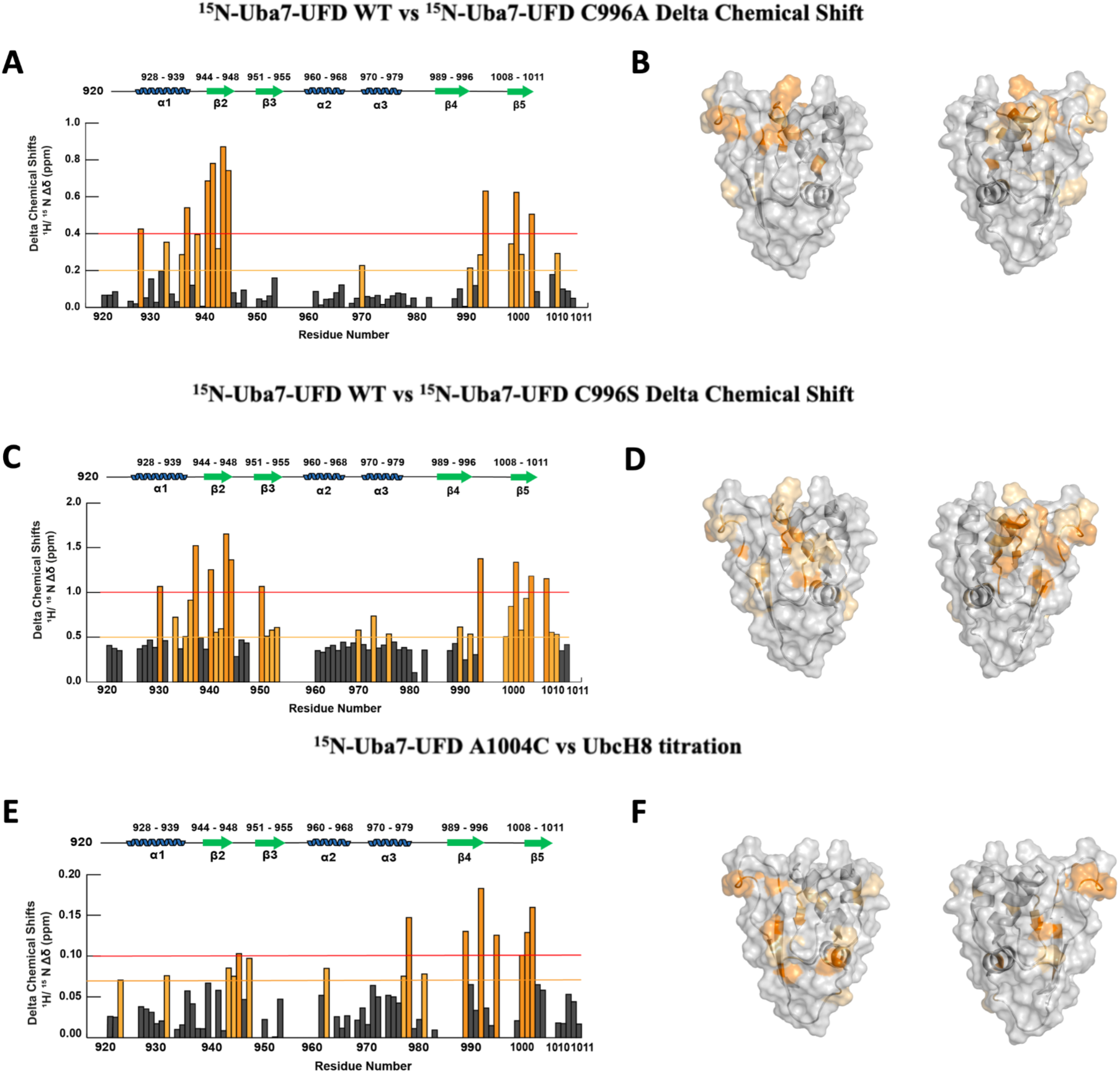
Investigation of the Effects of Mutations on ^1^H-^15^N Chemical Shift Values of Uba7-UFD A. ^15^N-Uba7-UFD WT vs ^15^N-Uba7-UFD C996A Delta Chemical Shift Map The residues colored orange possessed delta chemical shift larger than 0.4 ppm threshold (red line). The residues colored merigold possessed delta chemical shift larger than the 0.2 ppm threshold (merigold line). B.^15^N-Uba7-UFD WT vs ^15^N-Uba7-UFD C996A significant Delta Chemical Shifts Mapped on protein structure Residues with delta chemical shift larger than 0.4 ppm threshold (orange), residues with chemical shift perturbations larger than 0.2 ppm threshold (merigold) were mapped onto the Uba7-UFD structure C. ^15^N-Uba7-UFD WT vs ^15^N-Uba7-UFD C996S Delta Chemical Shift Map The residues colored orange possessed delta chemical shift larger than 1 ppm threshold (red line). The residues colored merigold possessed delta chemical shift larger than the 0.5 ppm threshold (merigold line). D. ^15^N-Uba7-UFD WT vs ^15^N-Uba7-UFD C996S significant Delta Chemical Shifts Mapped on protein structure Residues with delta chemical shift larger than 1 ppm threshold (orange), residues with chemical shift perturbations larger than 0.5 ppm threshold (merigold) were mapped onto the Uba7-UFD structure E. ^15^N-Uba7-UFD A1004C vs UbcH8 titration Chemical Shift Perturbation Map ^1^H-^15^N chemical shift perturbations for Uba7-UFD were calculated for each residue. The residues colored orange possessed chemical shift perturbations larger than 0.1 ppm threshold (red line). The residues colored merigold possessed chemical shift perturbations larger than the 0.06 ppm threshold (merigold line). F. ^15^N-Uba7-UFD A1004C significant Chemical Shift Perturbations Mapped on protein structure Residues with chemical shift perturbations larger than the 0.1 ppm threshold (orange), residues with chemical shift perturbations larger than 0.06 ppm threshold (merigold) and peak intensity changes that were larger than the threshold (blue) were mapped onto the Uba7-UFD structure

### NMR data driven docking of UbcH8 and Uba7-UFD complex

Next, we used the most substantial CSPs from our NMR titrations on the isolated domain interactions to restrain a model of the full-length Uba7-HFB/UbcH8 complex using the HADDOCK 2.4 NMR-based docking protocol (Vries et al., 2010). The AlphaFold Uba7 model (APSD: AF-P41226-F1-v4) and the UbcH8 crystal structure (PDB: 1WZV) were used as inputs and the resulting cluster with the lowest HADDOCK score was analyzed (Figure 4). The HADDOCK model illustrates how complexation is facilitated by two sets of charge-charge interactions; specifically, Uba7-UFD ^999^DDEDTA^1003^ and ^950^SALLYAAGWSPEK^962^ complement UbcH8 ^5^MRVVKE^10^ and ^25^LSSDD^29^, respectively. Furthermore, a series of hydrophobic and charged residues surrounding the active site cysteine undergoes conformational changes during the transthiolation stage (Figure 4). The spatial arrangement of these regions precisely orients UbcH8 to ensure efficient ISG transfer between catalytic cysteine residues. We also collected relaxation datasets for the Uba7-UFD:UbcH8 complex at a 1:2 molar ratio; however, several resonances disappeared upon complex formation, preventing reliable fitting of relaxation curves and thus precluding publication-quality analysis.

### Dynamics of Uba7 UFD

To characterize the internal motions and structural rigidity of the Uba7 UFD in solution, backbone ^15^N relaxation measurements were obtained, including longitudinal relaxation times (T_1_), transverse relaxation times (T_2_), T_1_/T_2_ ratios, and {^1^H}-^15^N heteronuclear NOE values (Figure 5). The overall relaxation profiles demonstrate that Uba7-UFD adopts a well-folded and dynamically stable β-grasp architecture, with relaxation parameters typical for a compact monomeric protein of this size (∼11 kDa). Regions corresponding to the β2 (residues 944-951) and β4 (residues 984-989) strands, which form the E2-binding surface are highlighted in orange. These segments show slightly lower {¹H}-¹⁵N NOE values and β2 segment show slightly higher T_2_ relaxation times, indicating enhanced local flexibility relative to the β-sheet core. Such mobility is often observed in loop regions or solvent-exposed β-strand edges that mediate transient interactions. The loop region spanning residues 996-1008 exhibits low {^1^H}-^15^N NOE values and prolonged T_2_ relaxation times, indicating enhanced backbone flexibility compared with the rest of the Uba7-UFD domain. This flexible segment forms part of the surface that interacts with ISG15 and likely contributes to its recognition. The observed mobility suggests that this loop can undergo transient conformational adjustments, allowing optimal positioning of ISG15 during E1-E2-substrate transfer. Such dynamic behavior may therefore play a crucial role in facilitating ISG15 conjugation and ensuring efficient catalysis within the ISGylation pathway.

To assess how E2 recruitment reshapes UFD dynamics, equivalent relaxation measurements were collected for ^15^N-labeled Uba7-UFD in the presence of unlabeled UbcH8 (Figure 5E–H). Consistent with complex formation, the relaxation behavior shifts toward values expected for a higher effective molecular size, including increased T_1_ and elevated T_1_/T_2_ ratios across the observable residues (Figure 5E,G). At the same time, residue-specific heterogeneity becomes more pronounced within the highlighted β2/β4 surface and adjacent segments, consistent with binding-coupled changes in local mobility and/or conformational exchange at the interface (Figure 5F,H). We note that complex formation can lead to the disappearance of a subset of resonances, which limits quantitative analysis for all residues and is consistent with intermediate-timescale exchange contributions in portions of the binding surface. The elevated mobility observed in β2/β4 and the 996–1008 loop provides a plausible physical basis for the disorder-to-order or adaptive-fit behavior inferred from binding and structural analyses, helping to reconcile how Uba7-UFD can remain stably folded while retaining the conformational plasticity required for efficient E2 recruitment and productive transthiolation during ISGylation.

Together, these data reveal that Uba7-UFD possesses a hierarchical dynamic profile, where structural rigidity within the β-sheet framework coexists with localized flexibility at the potential binding surface features likely critical for efficient recognition and alignment with its cognate E2 enzyme during ISGylation.

### Dynamics of E1 UFD binding surface on UbcH8

The dynamical behavior of the UbcH8 was also probed using ^15^N T1, ^15^N T2, and {^1^H}-^15^N heteronuclear NOE relaxation experiments (Lakomek, Ying and Bax, 2013). Figures 6A and 6B show the T1 and T2 relaxation rates plotted against the residue number and color mapped on the crystal structure of UbcH8 (PDB:1WZW), respectively. Residues with high T1 and low T2 relaxation rates corresponding to relatively rigid regions are colored in green whereas those displaying lower T1 and higher T2 relaxation rate hence higher flexibility are shown in yellow, orange, and red in the order of increasing flexibility. Gray color corresponds to the unassigned region. Figure 6C shows the heteronuclear ^15^N-^1^H NOE values plotted as a function of UbcH8 amino acid sequence. Fitting of data was done with the help of Bruker dynamics center and which showed more than 95% confidence level. All together, T1 and T2 relaxation rates and heteronuclear NOE values measured for UbcH8 consistently show that while the core of the protein is rigid, the outward-facing residues on the conserved N-terminal α-helix show enhanced flexibility, consistent with their proposed role in the UbcH8-binding interface (Figure 6D). Our docking studies show that this region of UbcH8 is directly involved in the interaction with Uba7-UFD. This relatively higher mobility may contribute to the E2’s interaction and function by facilitating conformational changes necessary for E1–E2 transthiolation.

### Conformational Flexibility of Uba7-UFD and Its Role in E2 Binding: Insights from Mutational Analysis

NMR solution structure of free Uba7-UFD indicated that the putative E2 binding region might be flexible and experiencing conformational plasticity, transitioning among multiple conformational states. Comparably, conformational flexibility/heterogeneity has been observed in E2 binding surfaces of SUMO (Wang et al., 2009) and Nedd8 E1 UFD (Elgin et al., 2012). Notably this conformational flexibility has been shown to be functionally important in SUMO conjugation (Wang et al., 2009).

To further explore this phenomenon, we designed and analyzed a series of mutants to determine how these alterations would affect E2 binding to Uba7-UFD. Specifically, we aimed to identify the role of individual amino acids in maintaining the structural integrity and interaction capabilities of the UFD domain. Given the high degree of plasticity observed in the putative E2 binding region, understanding how mutations influence this flexibility is crucial. This investigation not only provides insight into the specific amino acid interactions that are critical for binding but also sheds light on the overall mechanism by which Uba7-UFD achieves its conformational adaptability. By mapping these interactions and comparing the behavior of wild-type and mutant proteins, we sought to delineate the structural elements that enable effective binding and subsequent functional outcomes.

### Role C996 in Uba7-UFD conformational integrity and UbcH8 binding

In order to determine the effect of mutations on E2 binding to Uba7-UFD, seven mutants were created and five of them also titrated with UbcH8. Full length Uba7 protein contains a total of 17 cysteine residues, only one of which is in the UFD. Since this domain does not contain any other cysteine residue to form disulfide bridge with C996, this single cysteine probably remains reduced. C996 is located in the end of the second beta sheet, and while the subsequent amino acids show a very flexible behavior in the structure, cysteine and previous amino acids have formed a stable beta sheet secondary structure form.

We first mutated the free cysteine to alanine, which is the most preferred amino acid in mutation scans. Overlay of ^1^H-^15^N HSQC spectra of wild-type and C996A variant shows that the overall conformations of the mutant and wild-type proteins remain largely the same (Supp Figure 9). In contrast, changes in chemical shift values were observed around the mutated region, indicating a systematic reorganization in the preceding and subsequent beta sheets (Figure 7A). Figure 7B shows the residues affected by C996A mutation mapped onto the Uba7-UFD structure. Residues with chemical shift perturbations larger than 0.4 and 0.2 ppm colored in orange and merigold, respectively, are mostly clustered in neighboring B1, and B4 strands and alpha 1 helix. In order to determine the effect of mutation on E2 binding to Uba7-UFD, mutant protein was titrated with UbcH8. Supp Figure 10 shows the overlay of ^1^H-^15^N HSQC spectra of Uba7-UFD-C996A before and after addition of various amounts of UbcH8. No significant ^1^H-^15^N chemical shift perturbations in C996A mutant upon addition of UbcH8 was observed and addition of more and more of UbcH8 resulted in disappearance of some of the peaks involved in binding. These titration studies clearly indicate that C996 is important for Uba7-UFD-UbcH8 interaction (Supp Figure 10). Mutation of C996 to alanine appeared to have effect on UbcH8 binding. Since mutation of cysteine to alanine results in truncation of the side chain at the C-beta, the goal of C996S mutation was to replace –SH group with –OH group to see if this would restore the UbcH8 binding. The comparison of ^1^H-^15^N HSQC spectrum of C996S with that of wild-type shows that the overall conformation of C996S mutant is very similar to that of wild-type (Supp Figure 11) and as in C996A mutant, significant chemical shift perturbations are located around the mutation site (Figure 7C-7D). NMR binding studies carried out using ^15^N labeled Ubc7-UFD C996S further confirmed the role of C996 in UbcH8 binding since there were no significant chemical shift perturbations in the ^1^H-^15^N HSQC spectra of C996S mutant upon addition of UbcH8(Supp Figure 12). The failure of the -OH group in restoring the UbcH8 binding suggests that free -SH might not merely be involved in hydrogen bonding but might be important in providing necessary flexibility for UbcH8 interaction (Bhopatkar et al., 2020).

After determining that free cysteine was necessary for UbcH8 binding/for keeping Uba7-UFD in a suitable conformation for the UbcH8 binding we searched for amino acids in close proximity to C996 within Uba7-UFD to test the effect of breaking C996 intermolecular interactions on the conformation of the domain as well as on UbcH8 binding. Amino acids directly interacting with C996 were identified through NOE analysis. NOE peaks observed between C996 and R944 indicated that there is an interaction between these two amino acids. Therefore, an R944L mutant protein was generated, produced, and purified after refolding. Gel filtration chromatography of purified ^15^N-GBI-Uba7-UFD resulted in two separate peaks with the same molecular weight confirmed by SDS-PAGE analysis. Peaks were collected in separate fractions and analyzed by NMR. ^1^H-^15^N HSQC spectra showed that the peak eluted first from the gel filtration column was completely misfolded/aggregated (Supp Figure 13A) while the fraction corresponding to the second peak consisted of a mixture of both folded and misfolded fusion protein (Supp Figure 13B). Gel filtration chromatography and NMR results clearly show that R944 and its interaction with C996 are important for conformational stability of Uba7-UFD and mutation of R944 to alanine makes domain prominent to aggregation. The role of free cysteine amino acid in protein structure and internal dynamics is often overlooked. However, a study conducted by Mazmanian et al. in 2016 examined the participation and significance of free cysteine amino acids in hydrogen bonding networks in detail (Mazmanian et al., 2016). In the case of the Uba7-UFD complex, when a cysteine atom forms a hydrogen bond with arginine (R944), it exhibits distinct interaction patterns depending on the charge of the hydrogen bond donor. Specifically, cysteine forms shorter and more linear hydrogen bonds with positively charged residues such as histidine (His⁺), lysine (Lys⁺), and arginine (Arg⁺), where the hydrogen–sulfur distance (H···S) is less than 2.5 Å, and the bond angle (N−H···S) exceeds 150° (Mazmanian et al., 2016). In contrast, hydrogen bonds with neutral donors are longer (H···S > 2.5 Å) and less linear (N−H···S < 150°). Notably, Arg⁺ donates two hydrogen bonds to the cysteine thiol (CysSH), a characteristic not observed in other residues (Mazmanian et al., 2016). These findings align with previous studies demonstrating that positively charged hydrogen bond donors preferentially form stronger and more directional interactions with thiol groups, contributing to protein stability and conformational flexibility. Thus, the Uba7-UFD complex exemplifies the critical role of free cysteines in fine-tuning the stability and plasticity of protein structures.

### Effect of acidic loop flexibility and length on Uba7-UFD interaction with UbcH8

As mentioned above, our NMR solution structure of Uba7-UFD shows that the loop region that comes right after the C996 is relatively flexible. Since this loop is involved in interaction with UbcH8, one question is whether restricting this flexibility would prevent binding. To test this, another mutant was designed aiming to form a disulfide bond between the single cysteine 996 and alanine 1004. In this mutant, A1004 was individually replaced by cysteine. This variant was also expressed as inclusion bodies, solubilized with 7M GdnHCl and refolded by dialysis as described previously. Supp Figure 14 shows an overlay of ^1^H-^15^N HSQC spectra of A1004C mutant in the absence and presence of various amounts of UbcH8. Figure 7E shows the chemical shift perturbation analysis from the titration experiment. Unlike the previous mutations, this mutant variant still showed intrinsically interacted with UbcH8. As seen in Figure 7F, the interaction surface is also partially compatible with the interaction surface of the WT protein. Opposite to WT, A1004C mutant’s 1st beta sheet region did not show any interaction with UbcH8.

The acidic loop interacting with UbcH8 has been extended with a glycine amino acid as a final mutation approach (998GGG). The 998GGG mutant protein has been produced, purified, and subjected to ^1^H-^15^N HSQC spectrum collection. Although there were regional differences in the obtained spectrum, no significant changes were observed in the overall distribution (Supp Figure 15A). Unlabeled UbcH8 has been added to the obtained protein in a 1:1 ratio. Upon examination of the spectra obtained after mixing the apo form of 998GGG with UbcH8, no changes were observed in the chemical shift values indicating that integrity of the loop was necessary for UbcH8 binding. (Supp Figure 15B).

## Discussion

Ubiquitin and Ubiquitin-like post-translational modification systems are of great importance to organisms due to their extensive regulatory roles within cells. Elucidating the mechanisms at the atomic level within the enzymatic cascades that form the basis of Ub and Ubl post-translational modification systems has been a significant topic in biochemistry and structural biology for decades. Although the atomic-level details of the mechanisms of systems such as ubiquitination, neddylation, and sumoylation have been examined, structural studies on the ISGylation system were not available until recent years. Specifically, due to the high plasticity of the ubiquitin fold domain of the E1 enzyme, determining its structure has taken a considerable amount of time. During the preparation of our manuscript, two additional papers aimed at elucidating the details of ISGylation have been published (Wallace et al., 2023; Afsar et al., 2023). Although Cryo-EM and crystal structures are excellent tools for visualizing the overall mechanism of systems, they are still static and frozen structures. This makes them less effective for examining weak interaction surfaces and low-affinity complex structures. At this point, NMR spectroscopy continues to be an important structural biology method, providing atomic-level information to complement the remarkable findings from other methods and to elucidate the side effects caused by freezing. In the first part of our study, we determined the NMR structure of the ISGylation E1 enzyme and observed that the plasticity and unstable beta-sheet surfaces exhibited in solution are believed to play a significant role in the system’s operation and selectivity.

Contemporary Cryo-EM structures, especially those examining the E1-E2 interaction, present differing results regarding the amino acids and interaction angles involved in the interaction between the first alpha helix of the E2 enzyme and the high plasticity regions of the E1 UF domain (Wallace et al., 2023; Afsar et al., 2023). Despite the high overall resolutions of these structures, it is notable that the resolution in these domains sometimes drops regionally to 4-5 Å. The direct interactions obtained from our NMR data-driven model show greater alignment with the findings of Wallace et al. 2023.

Findings from T1 and T2 relaxation experiments, coupled with titration and docking data, provide a coherent picture suggesting that the N terminus region of Ubch8 is more flexible and that this flexibility is relevant to the interaction with the Uba7-UFD. NMR characterization studies of Uba7-UFD variants showed that both wild type and mutant domains have a high tendency to aggregate. None of the designed mutant Uba7-UFDs was more stable than the wild type Uba7-UFD. Our NMR mapping study showed that UbcH8 binding surface of Uba7-UFD is substantial and there are over 30 amino acids directly involved in binding. This study also showed that a single reduced cysteine residue at position 996 may play a major role in domain stability and fine tuning of plasticity.

## Conclusion

Our study has reiterated the importance of examining protein structures and dynamics in solution. It has been demonstrated that the UFD domain of the Uba7 protein undergoes conformational changes upon binding to UbcH8, consistent with the induced fit model. Structural analysis of the Uba7 UFD and complex structures obtained from other systems have shown that a large surface area of the UFD interacts with E2, highlighting the significance of this interaction in E1-E2 binding and proper E2 selection. Additionally, structural investigations have revealed that a region on the E1 UFD interacts with Ubiquitin/Ubiquitin-like proteins. It is important to conduct new studies to demonstrate the importance of this interaction in structural and functional regulation.

## Author Contributions

**Çağdaş Dağ:** Conceptualization, Methodology, Supervision Funding acquisition, Investigation, Formal analysis, Writing- Reviewing and Editing, Writing-Original draft preparation **Alp Eren Kazar:**Investigation, Formal analysis, Visualization, Writing- Reviewing and Editing, Writing-Original draft preparation **Emine Sonay Elgin:** Conceptualization, Methodology, Supervision Funding acquisition, Investigation, Formal analysis, Writing- Reviewing and Editing, Writing-Original draft preparation **Arthur L. Haas:** Conceptualization, Supervision **Kerem Kahraman**: Investigation, Formal analysis, Visualization. **Oktay Göcenler:** Formal analysis, Investigation Writing - Original Draft, Visualization **Mahil Lambert:** Formal analysis, Investigation, Writing - Original Draft, Visualization **Volker Dötsch**: Conceptualization, Supervision, Writing-Reviewing and Editing **Cansu D. Tozkoparan:**Visualization, Formal analysis **Joshua J. Ziarek:** Conceptualization, Writing- Reviewing and Editing

## ACKNOWLEDGMENT

We gratefully acknowledge Prof. Brian F. Volkman (Medical College of Wisconsin) for generously opening his laboratory to us and providing access to critical instrumentation and consumables (GM64598-03). His support and willingness to share resources were invaluable for the successful execution of this work. We sincerely thank him for his scientific generosity and hospitality. ESE acknowledges support from TÜBİTAK (Project No: 104T193). CD acknowledges support from TÜBİTAK (Project No: 120Z594, 122Z747). JJZ acknowledges support from National Institutes of Health grant R35GM143054. The authors acknowledge the use of the services and facilities of n^2^STAR-Koç University Nanofabrication and Nanocharacterization Center for Scientific and Technological Advanced Research. This study made use of the National Magnetic Resonance Facility at Madison, which is supported by NIH grant R24GM141526 and P41GM103399. This work benefited from access to [CERM and BMRZ] and has been supported by iNEXT-Discovery, project number 871037, funded by the Horizon 2020 program of the European Commission. Financial support by the Access to Research Infrastructures activity in the 7th Framework Programme of the EC (Project number: 261863, Bio-NMR) for conducting the research is gratefully acknowledged. This study made use of NMRbox: National Center for Biomolecular NMR Data Processing and Analysis, a Biomedical Technology Research Resource (BTRR), which is supported by NIH grant P41GM111135 (NIGMS).

---

**Supplementary Figure 1.**
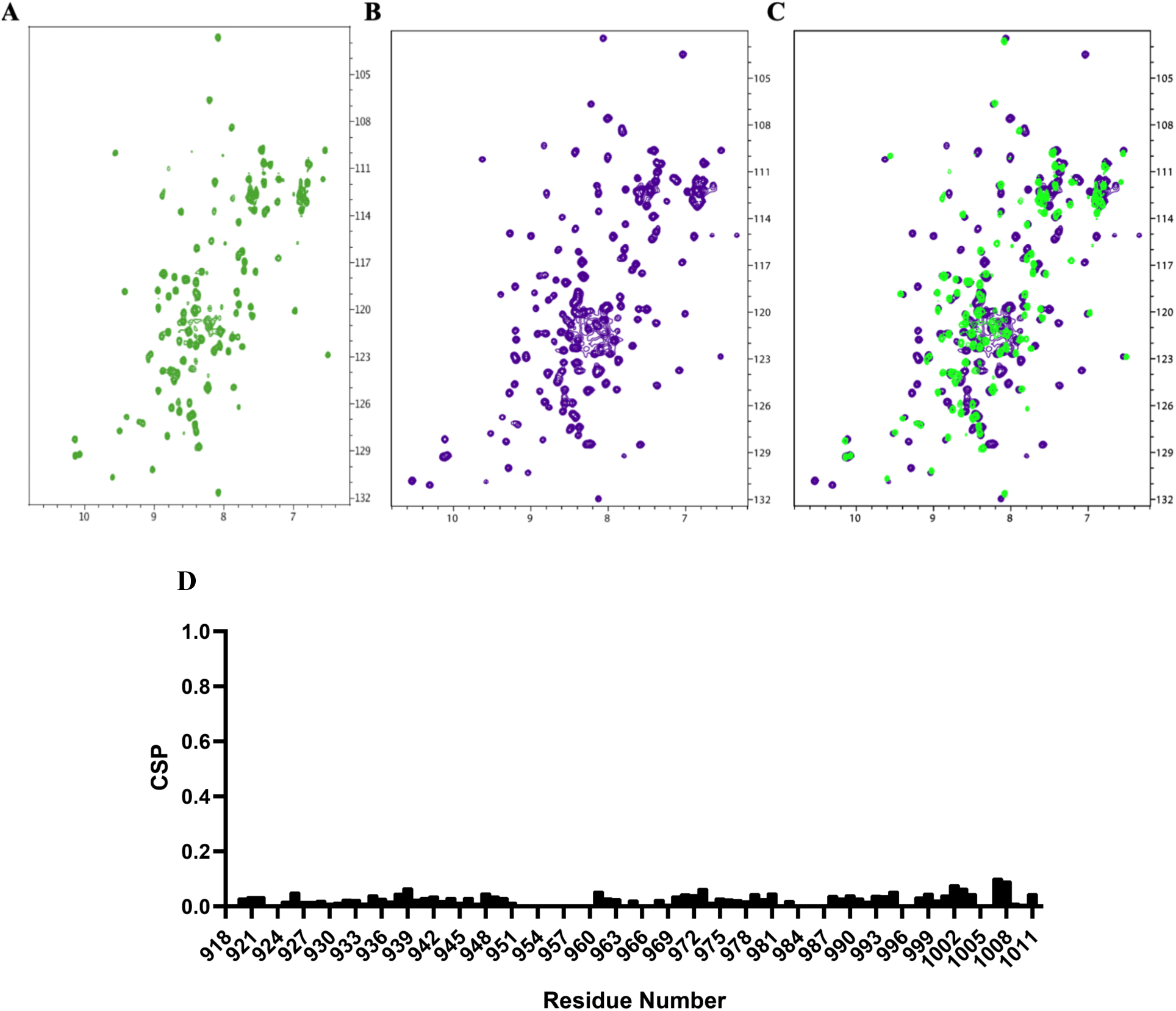
Human Uba7 enzyme ubiquitin fold domain ^1^H-^15^N HSQC spectrum A. ^1^H-^15^N HSQC spectrum of Uba7-UFD after GBI fusion protein cut. B. ^1^H-^15^N HSQC spectrum of GBI-Uba7-UFD C. ^1^H-^15^N HSQC spectrum of GBI-Uba7-UFD overlaid with ^1^H-^15^N HSQC spectrum of Uba7-UFD after GBI fusion protein cut D. Chemical shift perturbations (CSPs) analysis of Uba7-UFD with and without the GB1 fusion tag.

**Supplementary Figure 2.**
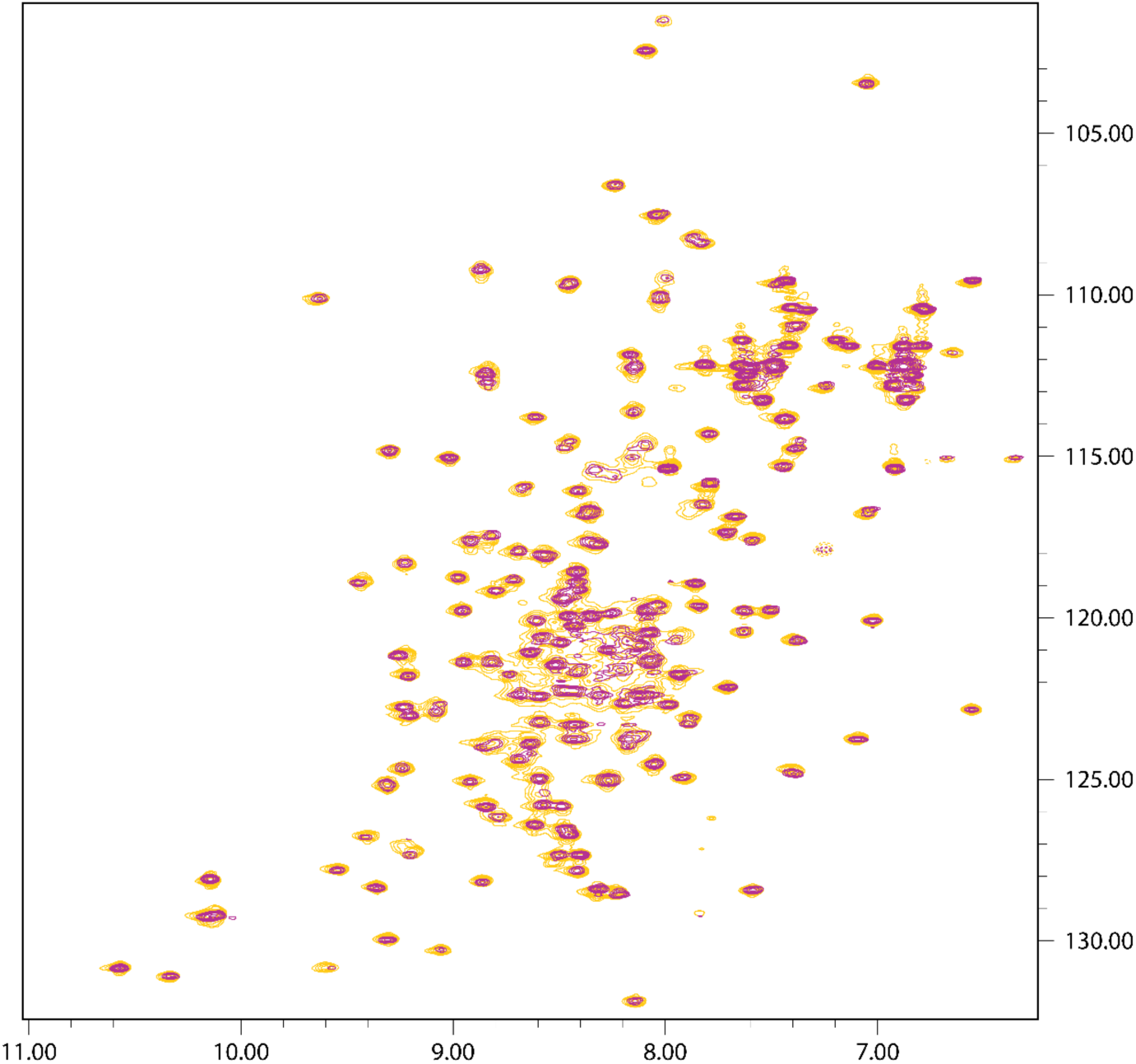
^1^H-^15^N HSQC spectrum of 0.4 mM ^15^N-GBI-Uba7-UFD (wt) protein (yellow on day 1, red on day 9)

**Supplementary Figure 3.**
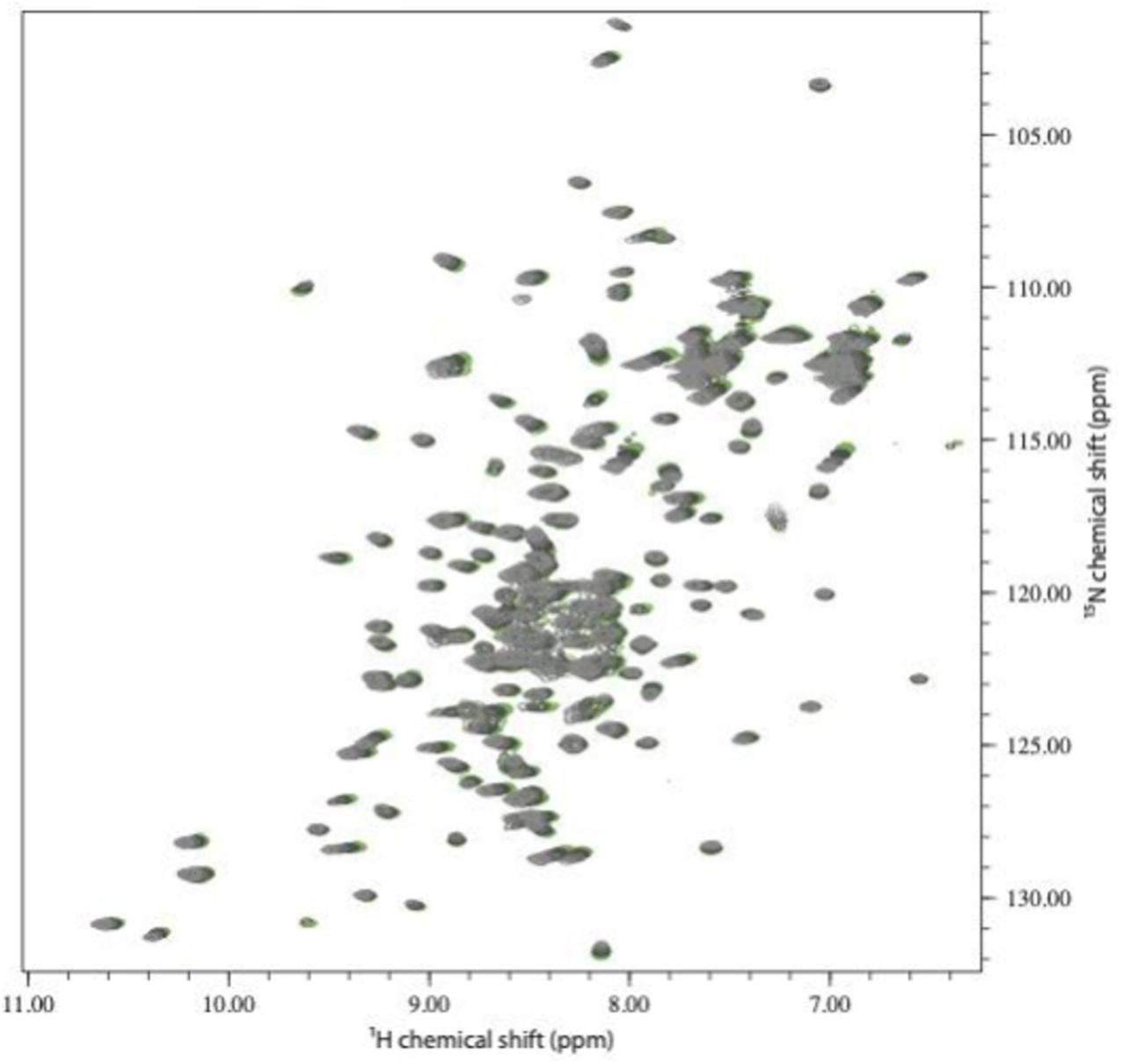
Overlay of ^15^N-GBI-Uba7-UFD ^1^H-^15^N HSQC spectra at different temperatures. (green 25°C, black 20°C, dark gray 15°C, light gray 10°C)

**Supplementary Figure 4.**
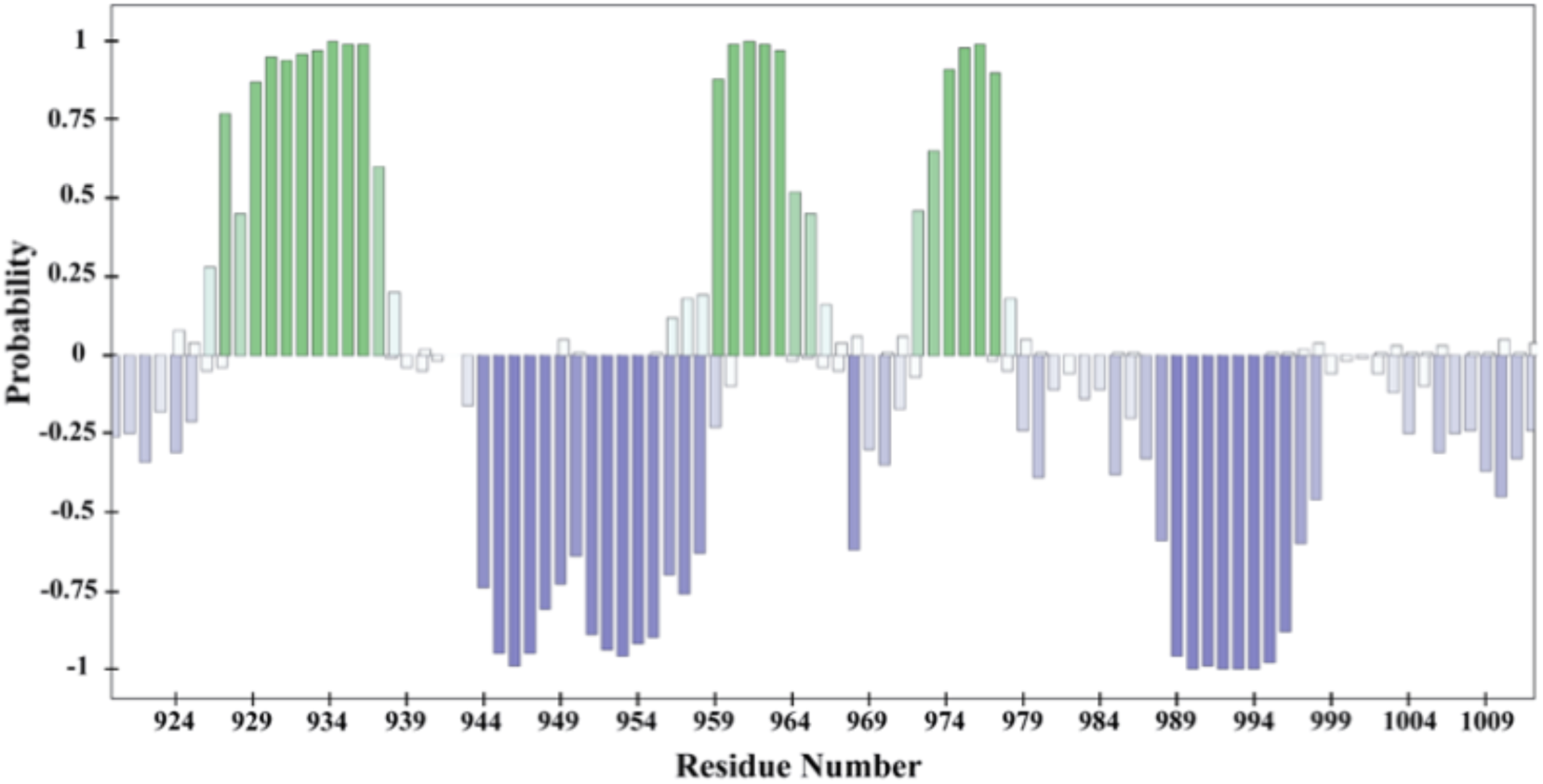
NMR chemical-shift-based TALOS-N secondary-structure prediction of Uba7-UFD

**Supplementary Figure 5.**
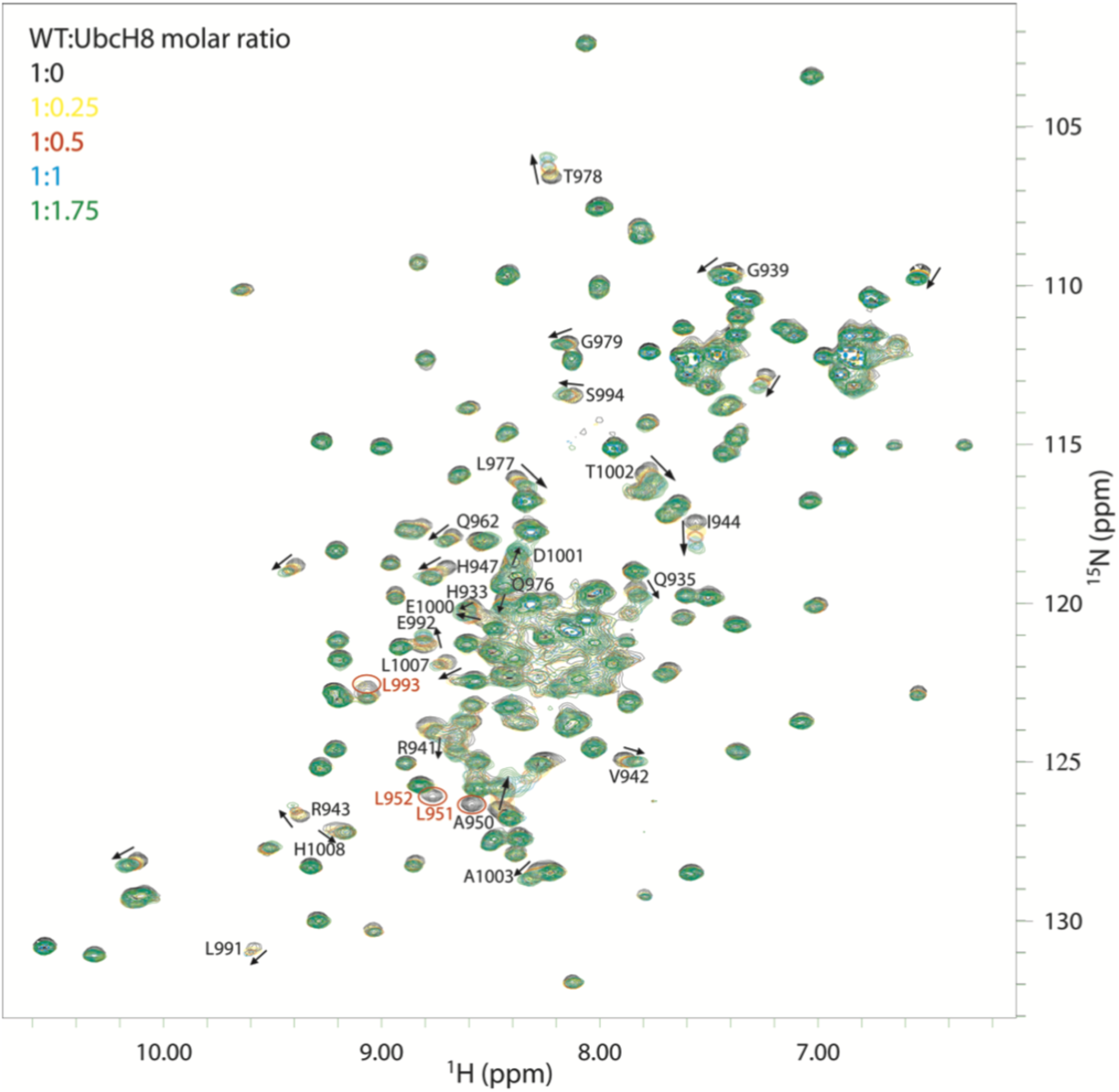
Superposition of the ^1^H-^15^N HSQC spectra of free (black) and various amounts of UbcH8 bound forms of wild type ^15^N-GBI-Uba7-UFD. GBI-Uba7-UFD:UbcH8 molar ratios in the bound forms are indicated in yellow for 1:0.25, red for 1:0.5, blue for 1:1, and green for 1:1.75. Peaks that disappear upon addition of 1:0.25 molar ratio of UbcH8 are indicated with red circles. Residues that show combined chemical shift perturbations of larger than 0.2 are indicated with an arrow and black one letter code followed by the residue number.

**Supplementary Figure 6.**
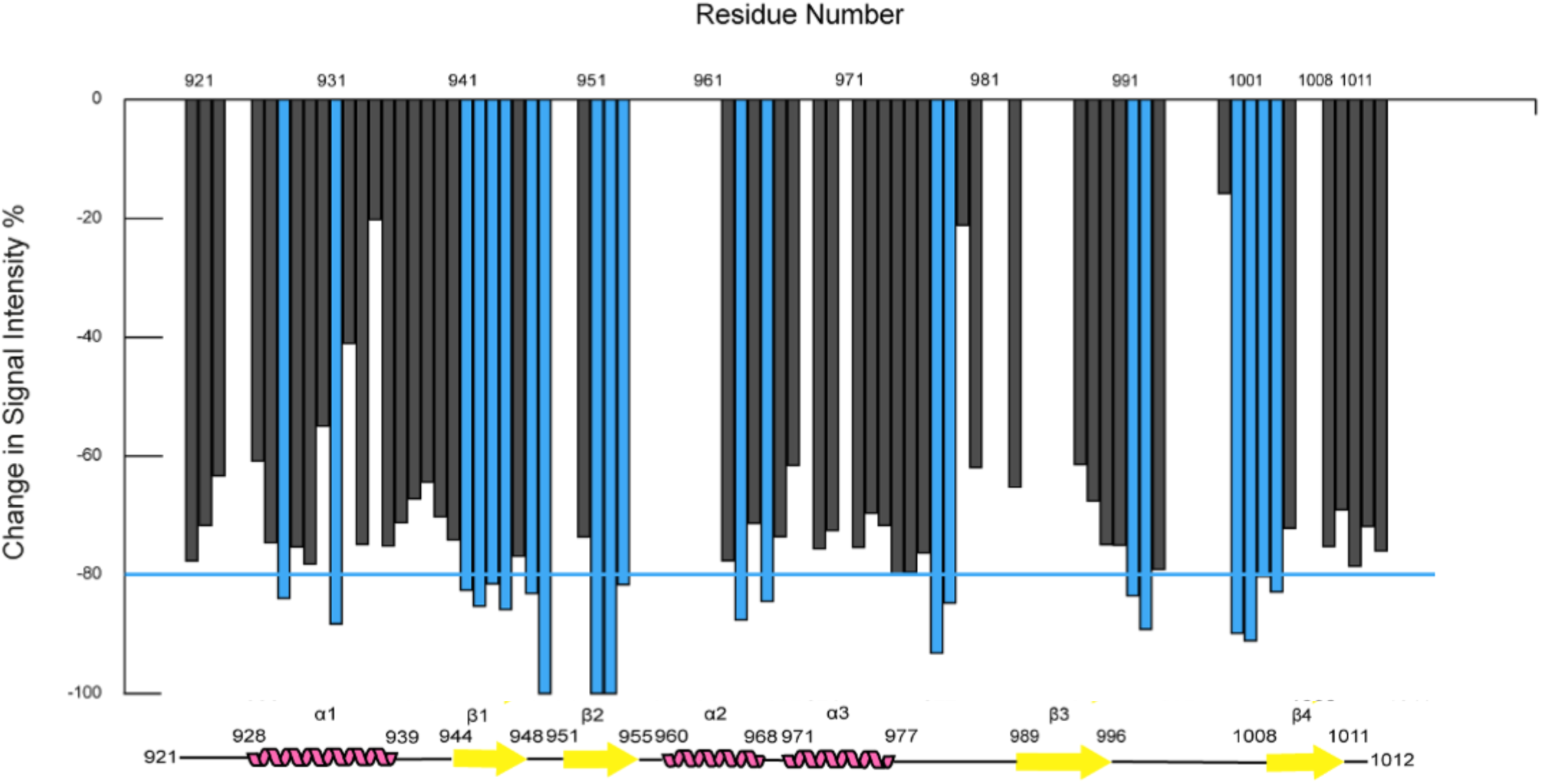
^15^N-Uba7-UFD vs UbcH8 titration Delta Chemical Shift Intensity Graph ^1^H-^15^N HSQC peak intensity changes for Uba7-UFD were calculated for each residue. The residues colored blue possessed peak intensity changes that were larger than the threshold (blue line)

**Supplementary Figure 7.**
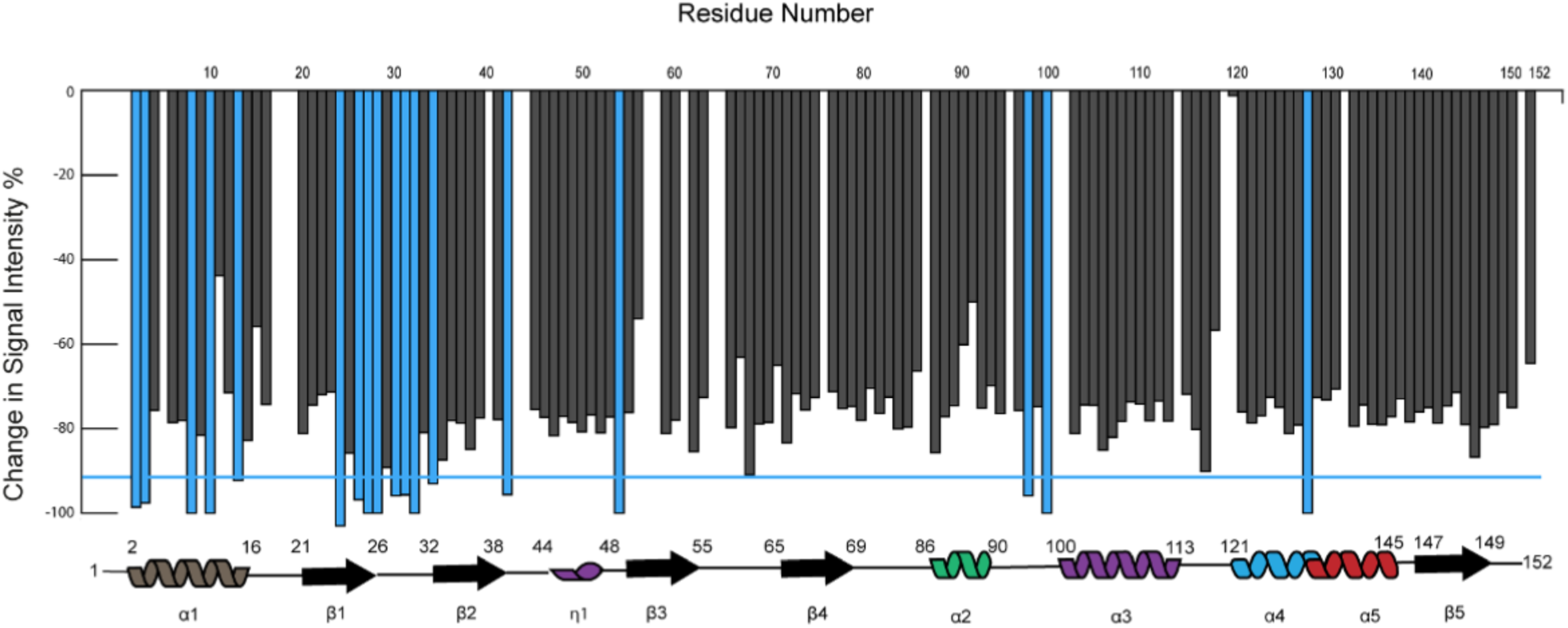
^15^N-UbcH8 vs Uba7-UFD titration Delta Chemical Shift Intensity Graph ^1^H-^15^N HSQC peak intensity changes for UbcH8 were calculated for each residue. The residues colored blue possessed peak intensity changes that were larger than the threshold (blue line)

**Supplementary Figure 8.**
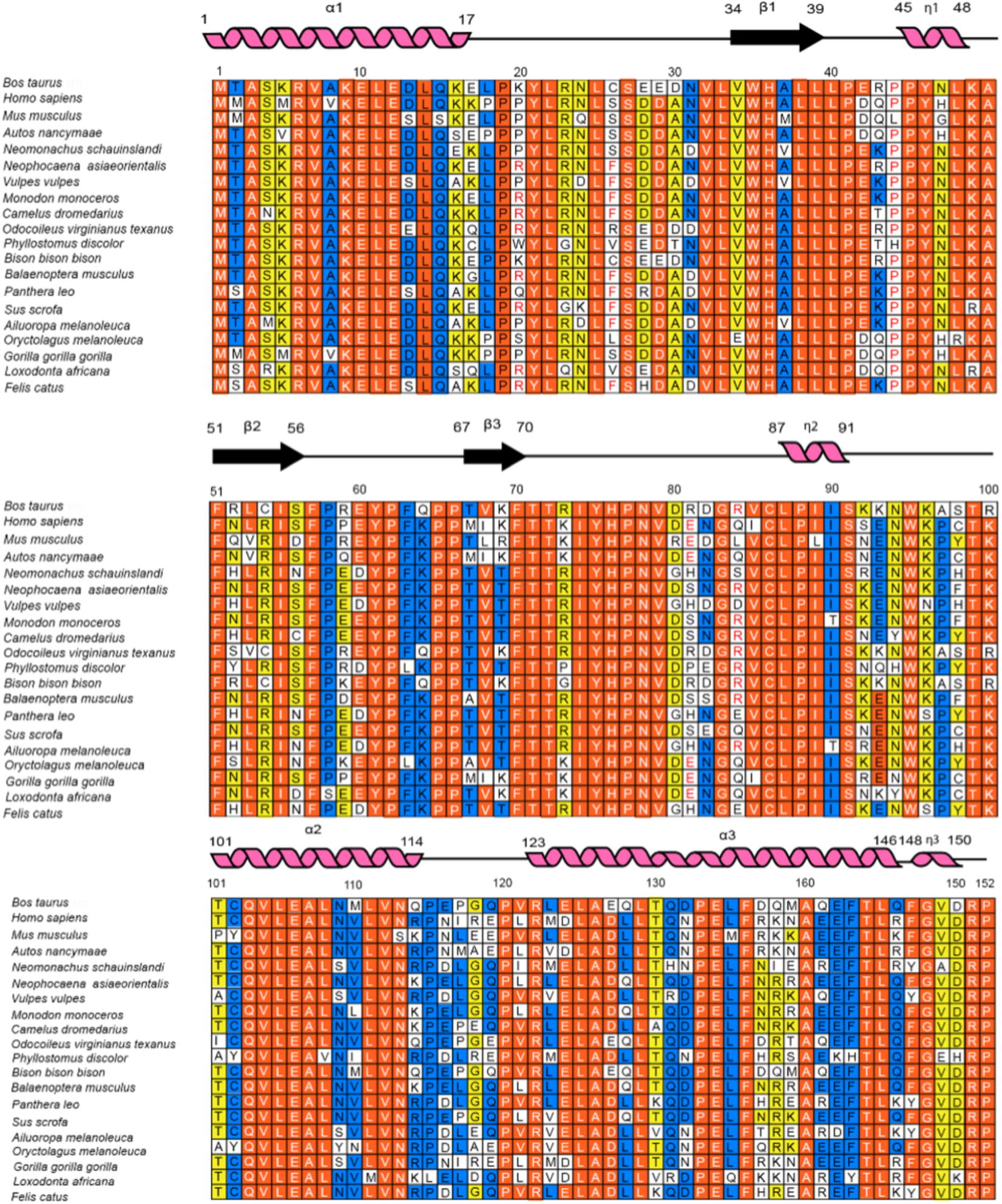
Sequence alignment of UbcH8 from different organisms.

**Supplementary Figure 9.**
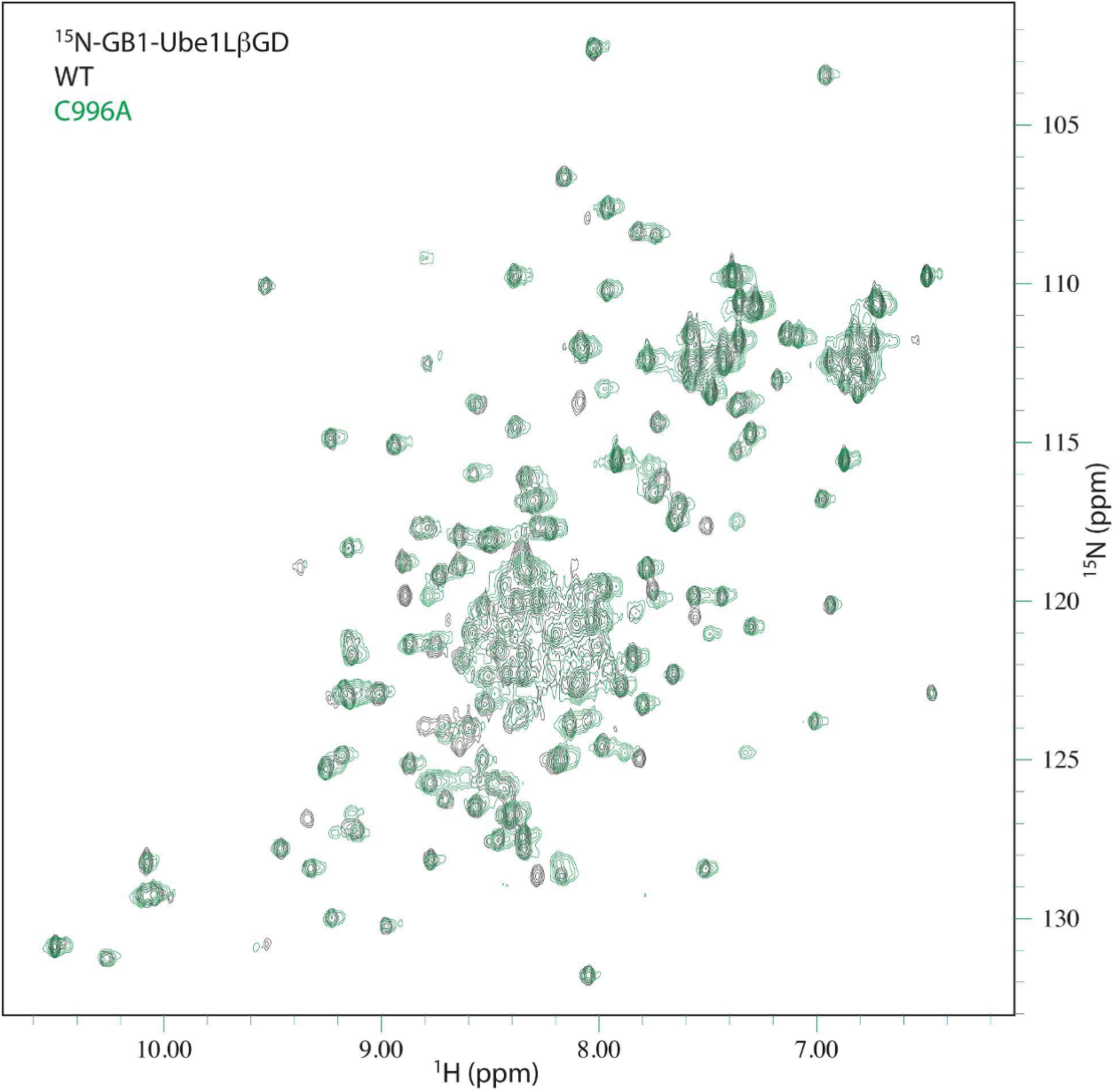
Superposition of 2D ^1^H-^15^N HSQCs of wt ^15^N-GBI-Uba7-UFD (black) with, C996A mutant (green).

**Supplementary Figure 10.**
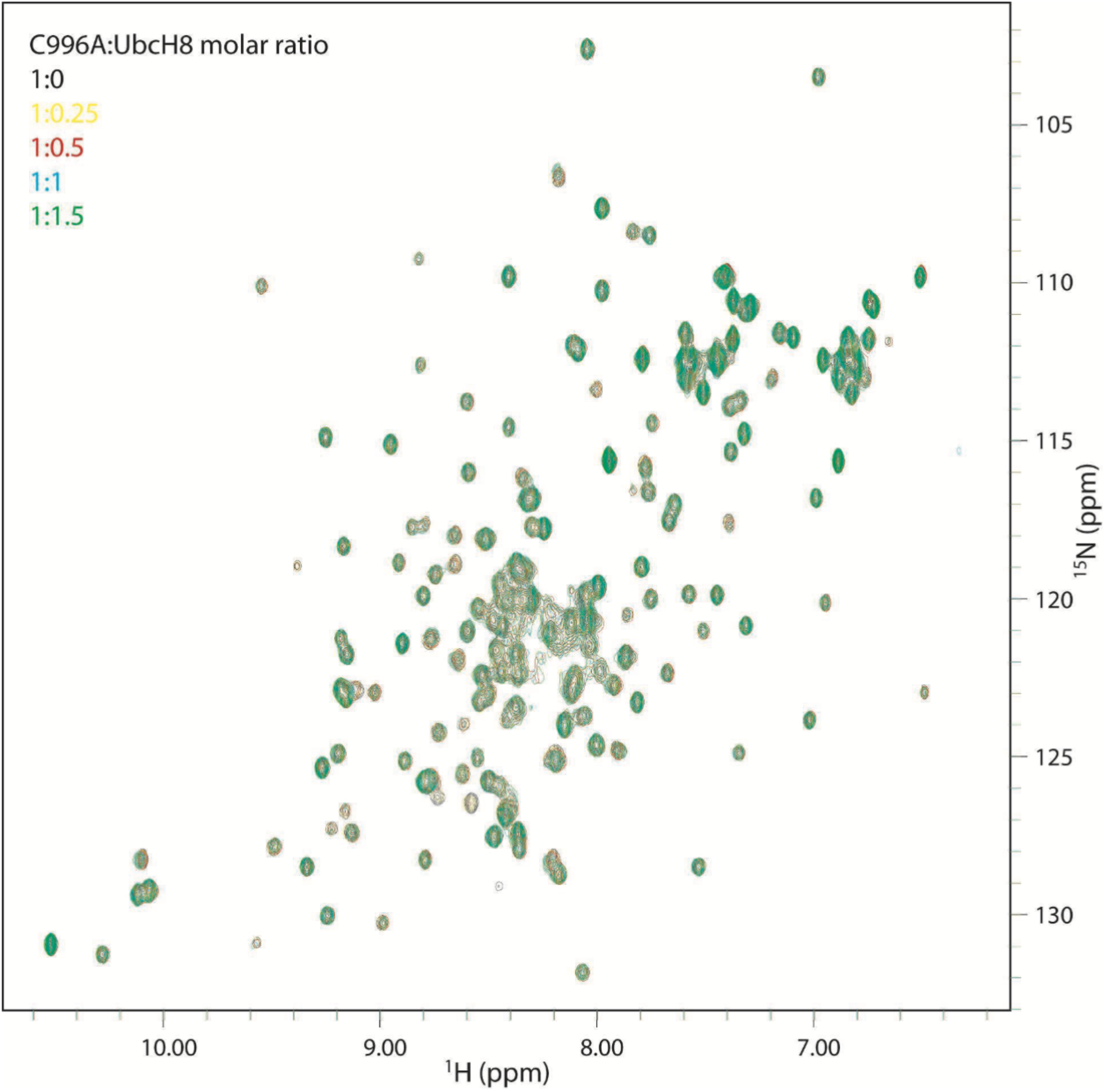
Superposition of the ^1^H-^15^N HSQC spectra of free (black) and various amounts of UbcH8 bound forms of ^15^N-GBI-Uba7-UFD, C996A mutant. GBI-Uba7-UFD:UbcH8 molar ratios in the bound forms are indicated in yellow for 1:0.25, red for 1:0.5, blue for 1:1, and green for 1:1.5.

**Supplementary Figure 11.**
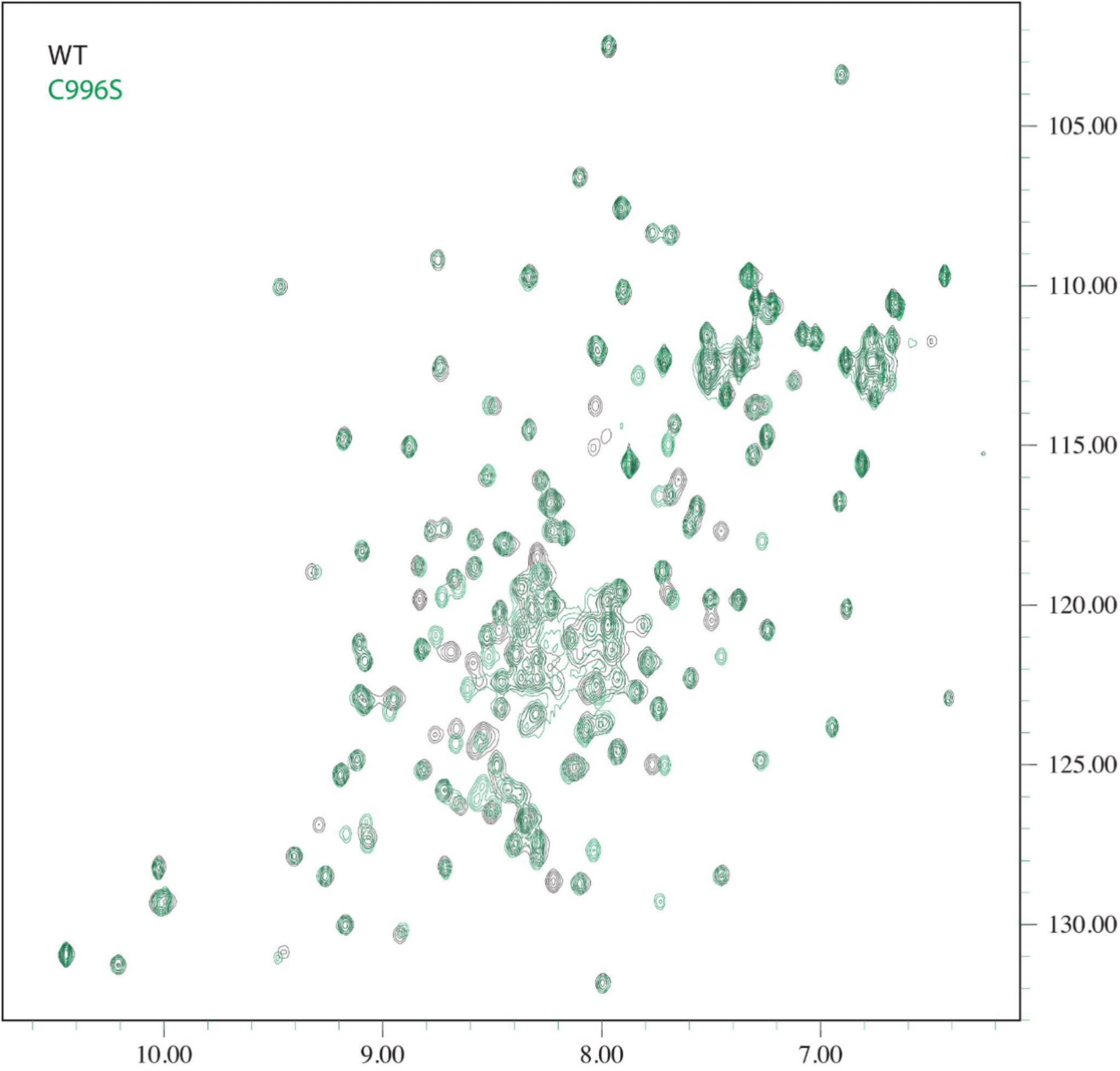
Superposition of 2D ^1^H-^15^N HSQCs of wt ^15^N-GBI-Uba7-UFD (black) with, C996S mutant (green).

**Supplementary Figure 12.**
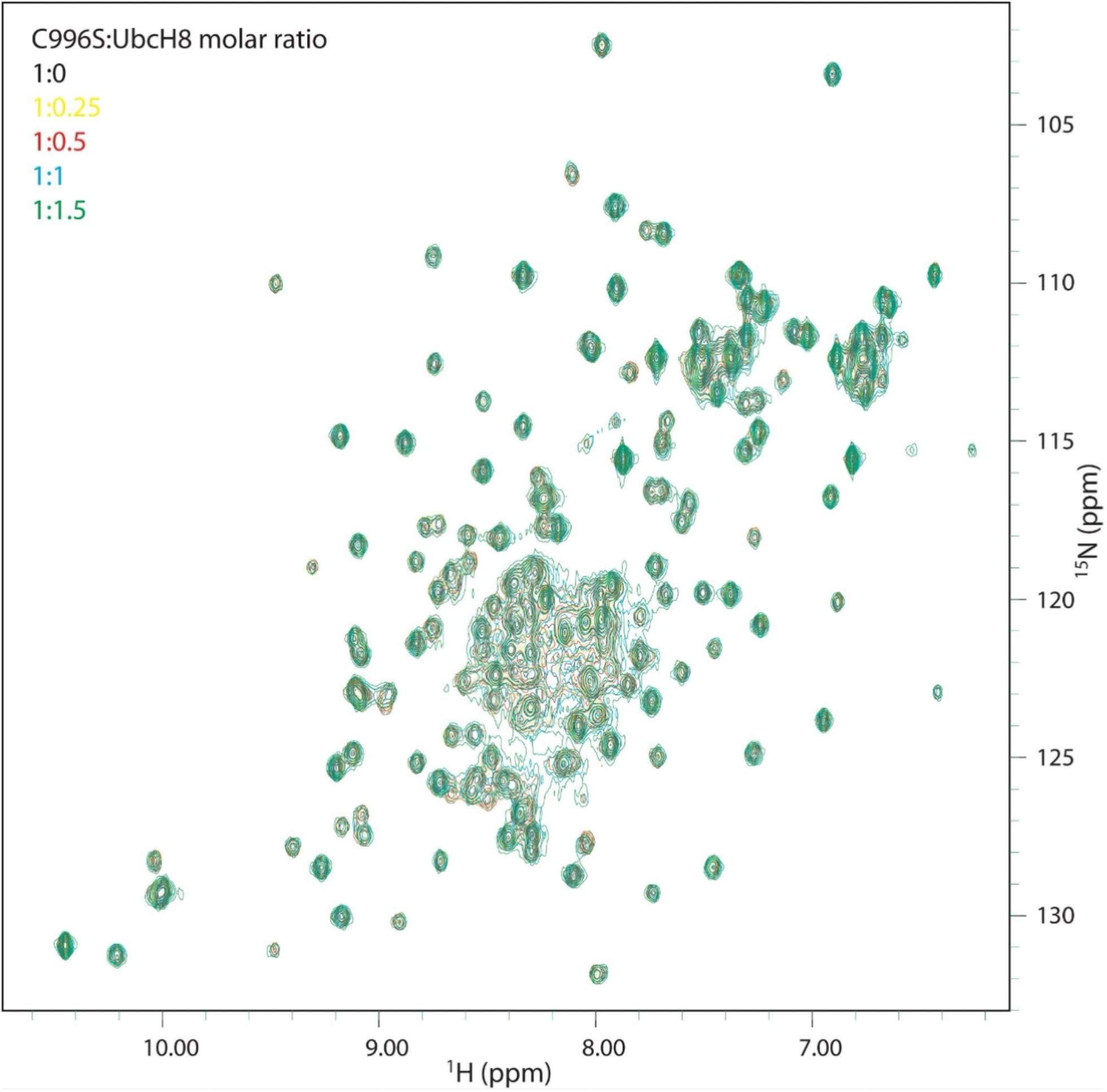
Superposition of the ^1^H-^15^N HSQC spectra of free (black) and various amounts of UbcH8 bound forms of ^15^N-GBI-Uba7-UFD, C996S mutant. GBI-Uba7-UFD:UbcH8 molar ratios in the bound forms are indicated in yellow for 1:0.25, red for 1:0.5, blue for 1:1, and green for 1:1.5.

**Supplementary Figure 13.**
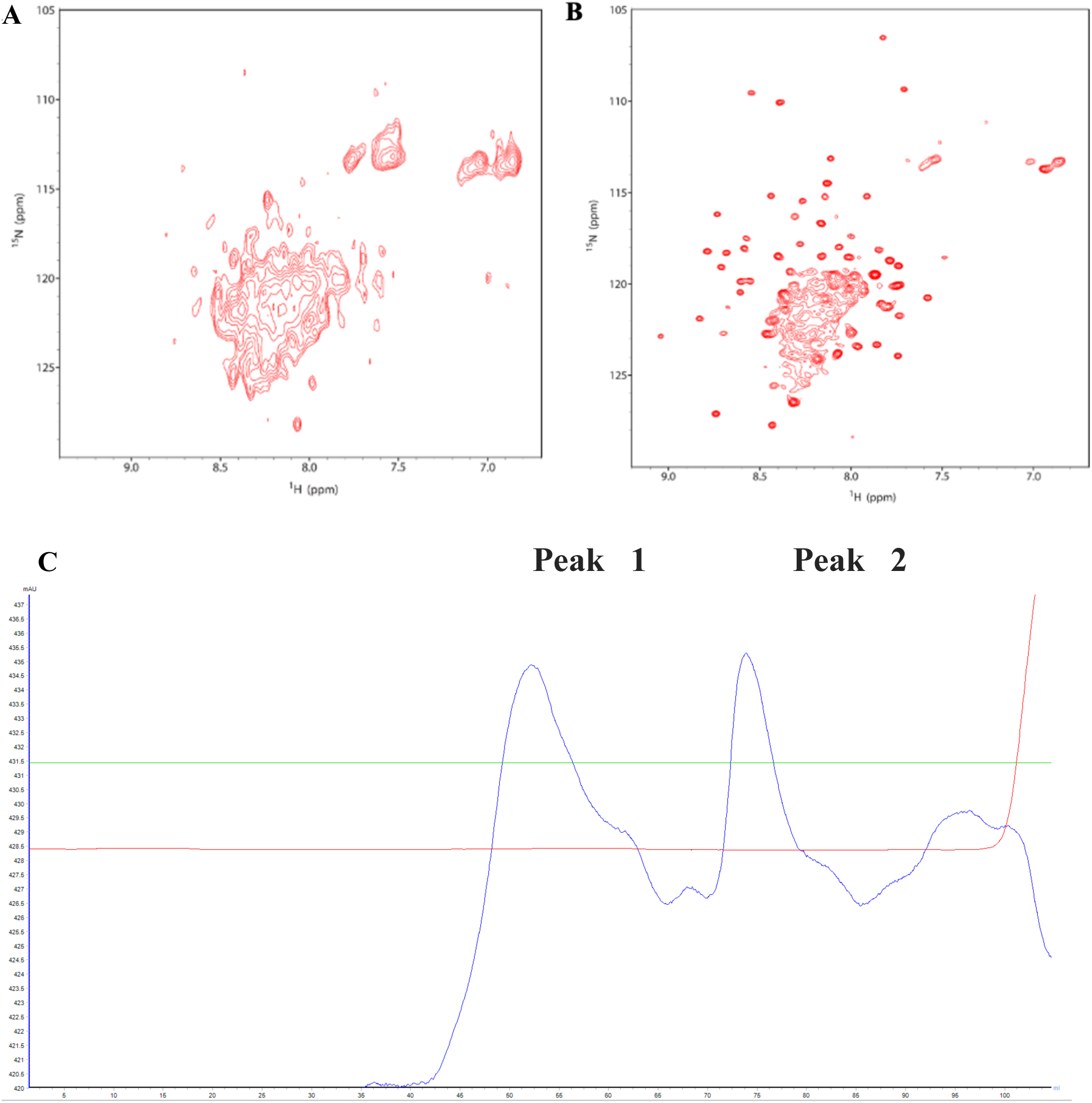
^1^H-^15^N HSQC spectra of free form of R944L mutant ^15^N-GBI-Uba7-UFD. A. ^1^H-^15^N HSQC spectrum from first gel filtration peak. B. ^1^H-^15^N HSQC spectrum from second gel filtration peak. C. S75 GF chromatogram

**Supplementary Figure 14.**
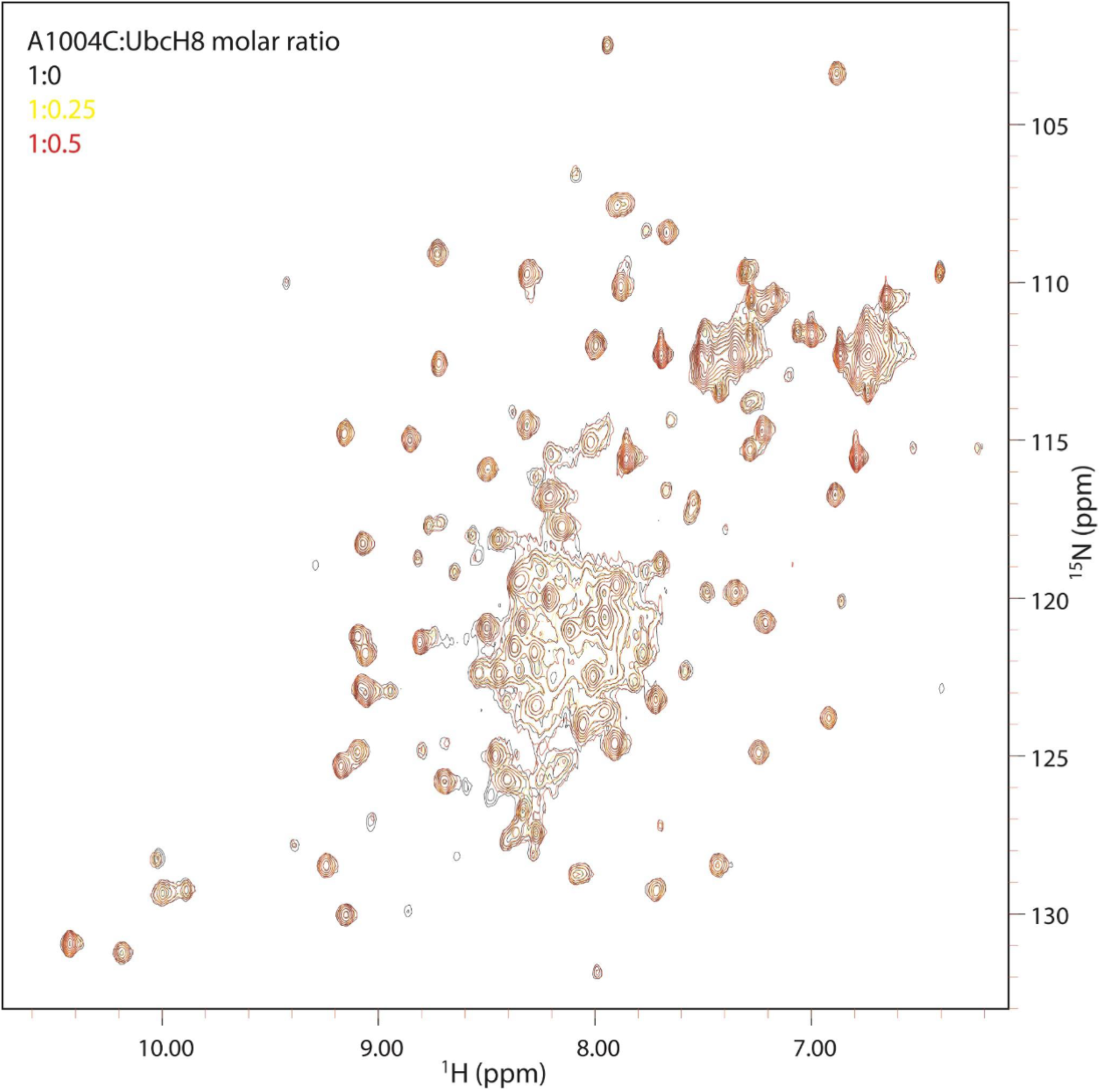
Superposition of the ^1^H-^15^N HSQC spectra of free (black) and various amounts of UbcH8 bound forms of ^15^N-GBI-Uba7-UFD, A1004C mutant. GBI-Uba7-UFD:UbcH8 molar ratios in the bound forms are indicated in yellow for 1:0.25, red for 1:0.5.

**Supplementary Figure 15.**
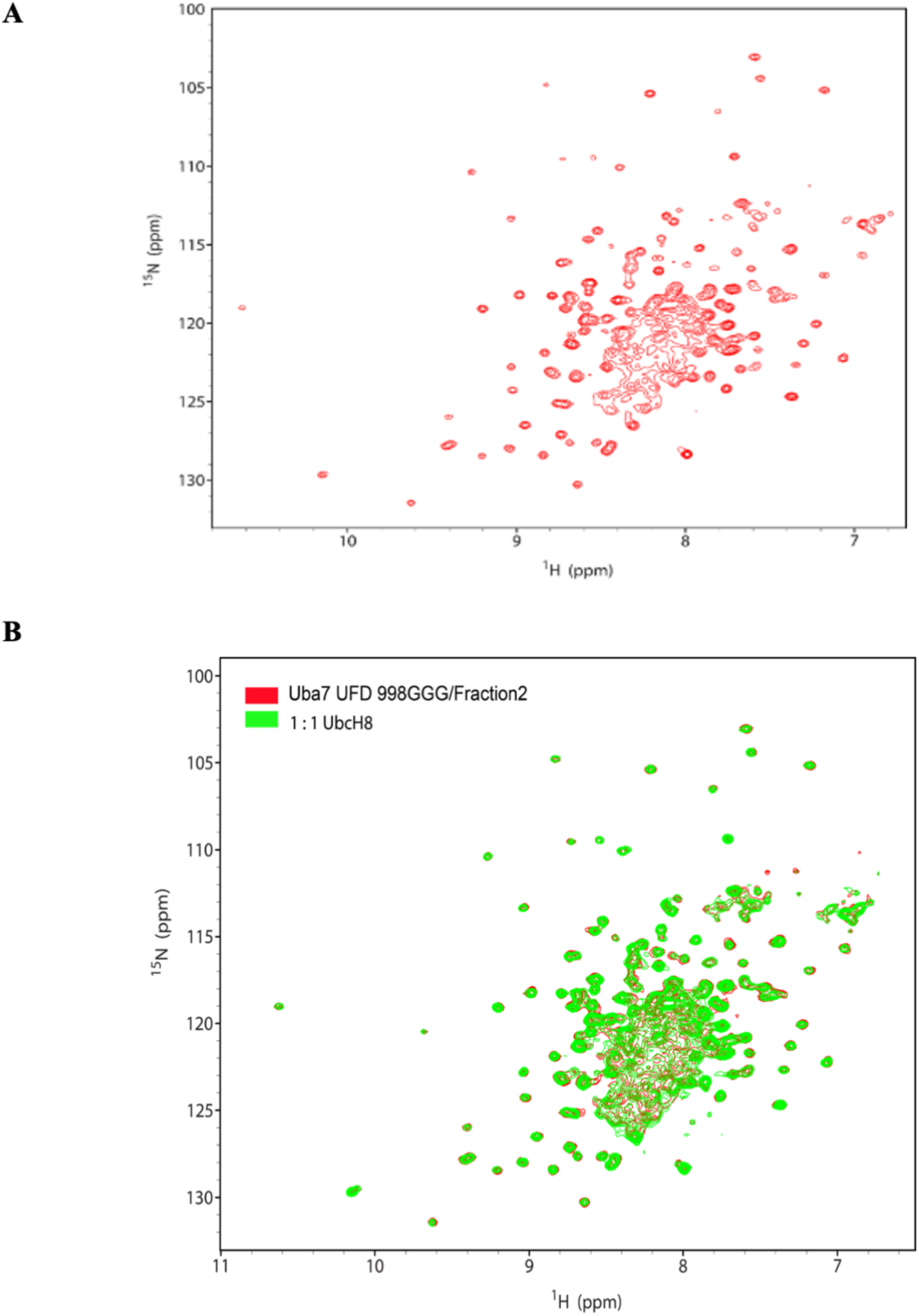
2D NMR analysis of Uba7-UFD 998GGG mutant A. ^1^H-^15^N HSQC spectra of free form of 998GGG mutant ^15^N-GBI-Uba7-UFD B. Superposition of the ^1^H-^15^N HSQC spectra of free (red) and 1:1 ratio of UbcH8 bound form of ^15^N-GBI-Uba7-UFD, 998GGG mutant (Green).

